# A single dose of cocaine rewires the 3D genome structure of midbrain dopamine neurons

**DOI:** 10.1101/2024.05.10.593308

**Authors:** Dominik Szabó, Vedran Franke, Simona Bianco, Mykhailo Y. Batiuk, Eleanor J. Paul, Alexander Kukalev, Ulrich G. Pfisterer, Ibai Irastorza-Azcarate, Andrea M. Chiariello, Samuel Demharter, Luna Zea-Redondo, Jose P. Lopez-Atalaya, Mario Nicodemi, Altuna Akalin, Konstantin Khodosevich, Mark A. Ungless, Warren Winick-Ng, Ana Pombo

**Author notes:** These authors contributed equally to this work (Dominik Szabó, Vedran Franke). Centre for Developmental Neurobiology, Institute of Psychiatry, Psychology and Neuroscience, King’s College London, London SE1 1UL, UK. Medical Research Council Centre for Neurodevelopmental Disorders, King’s College London, London SE1 1UL, UK. Laboratory of Neuroepigenetics, Brain Mind Institute, School of Life Sciences, École Polytechnique Fédérale Lausanne, Lausanne, 1015 Switzerland (Mykhailo Y. Batiuk). Center for Translational Genomics, Lund University, Lund, 221 85 Sweden (Ulrich G. Pfisterer). These authors jointly supervised this work (Warren Winick-Ng, Ana Pombo).

## Abstract

Midbrain dopamine neurons (DNs) respond to a first exposure to addictive drugs and play key roles in chronic drug usage^1–3^. As the synaptic and transcriptional changes that follow an acute cocaine exposure are mostly resolved within a few days^4,5^, the molecular changes that encode the long-term cellular memory of the exposure within DNs remain unknown. To investigate whether a single cocaine exposure induces long-term changes in the 3D genome structure of DNs, we applied Genome Architecture Mapping and single nucleus transcriptomic analyses in the mouse midbrain. We found extensive rewiring of 3D genome architecture at 24 hours past exposure which remains or worsens by 14 days, outlasting transcriptional responses. The cocaine-induced chromatin rewiring occurs at all genomic scales and affects genes with major roles in cocaine-induced synaptic changes. A single cocaine exposure triggers extensive long-lasting changes in chromatin condensation in post-synaptic and post-transcriptional regulatory genes, for example the unfolding of *Rbfox1* which becomes most prominent 14 days post exposure. Finally, structurally remodeled genes are most expressed in a specific DN sub-type characterized by low expression of the dopamine auto-receptor *Drd2*, a key feature of highly cocaine-sensitive cells. These results reveal an important role for long-lasting 3D genome remodelling in the cellular memory of a single cocaine exposure, providing new hypotheses for understanding the inception of drug addiction and 3D genome plasticity.

## Main

How is an initial exposure to addictive drugs encoded in cellular memory? Dopaminergic neurons (DNs) are critical players in the first response to drug-associated reward learning and reinforcement; a single exposure to cocaine induces long-term potentiation (LTP) of DN synapses in the midbrain ventral tegmental region (VTA), lasting up to 10 days^1^. These long-term effects of an initial exposure to addictive drugs, or other LTP activation events, are independent of long-lasting changes in gene expression, which are reported to occur and be resolved within the first 6-24 hours^4–7^. The first drug exposure is thought to alter the state of DNs, priming them for a much stronger and persistent, memory-associated, LTP induction that occurs after a second or multiple doses^8^, reflecting long-term responses seen in models of addiction learning paradigms^9^. However, without detectable lasting transcriptional or electrophysiological changes, it remains unknown how the cellular memory of a single drug exposure is encoded in DNs.

Our prior work mapping 3D genome structures in the adult mouse brain showed that addiction genes establish specific chromatin structures in VTA DNs, that are absent in other brain cell types^10^. Recent reports show that multiple cocaine exposures alter chromatin organization and the epigenetic state of addiction loci in brain regions involved in secondary cocaine-responses and which receive input from VTA DNs^11,12^. Altered chromatin looping has also been reported *in vitro*, after chronic pharmacological activation of glutamate neurons for 1 day, at immediate-early genes (IEGs)^13^. Regardless of all current efforts to understand the onset of addiction^14^, it remains unknown whether a single exposure to an addictive drug is sufficient to induce changes in chromatin structure *in vivo* in VTA DNs, whether and which chromatin changes are long lasting, and which genes are affected. In this study, we show that the 3D genome structure of VTA DNs is extensively rewired 24 hours after a single exposure to cocaine, with structural changes that can last or become more extensive after 14 days, a time-frame that is well beyond the resolution of transient transcriptional^4,5^ and LTP-associated^1^ effects. Our work implicates the remodelling of 3D genome structure as a mechanism for the cellular memory of a first drug exposure and as a basis for the onset of addiction, and identifies susceptible genes.

### Mapping the 3D genome structure and transcriptome of midbrain dopamine neurons upon a single cocaine exposure

To study the short- and long-term effects of a single exposure to cocaine on 3D genome structure and gene expression, we injected mice with either cocaine (15 mg/kg)^1,15^ or saline, and isolated DNs from the midbrain VTA after 1 or 14 days (in total 26 adult mice; **Fig. 1a**). We chose 1 day post-exposure to identify rewired 3D genomic structures that would persist beyond the timeframe of previously reported transcriptional changes^4,5^, and also analyzed changes at 14 days to discover long-lasting effects well past reported LTP responses^1,16^.

**Figure 1.**
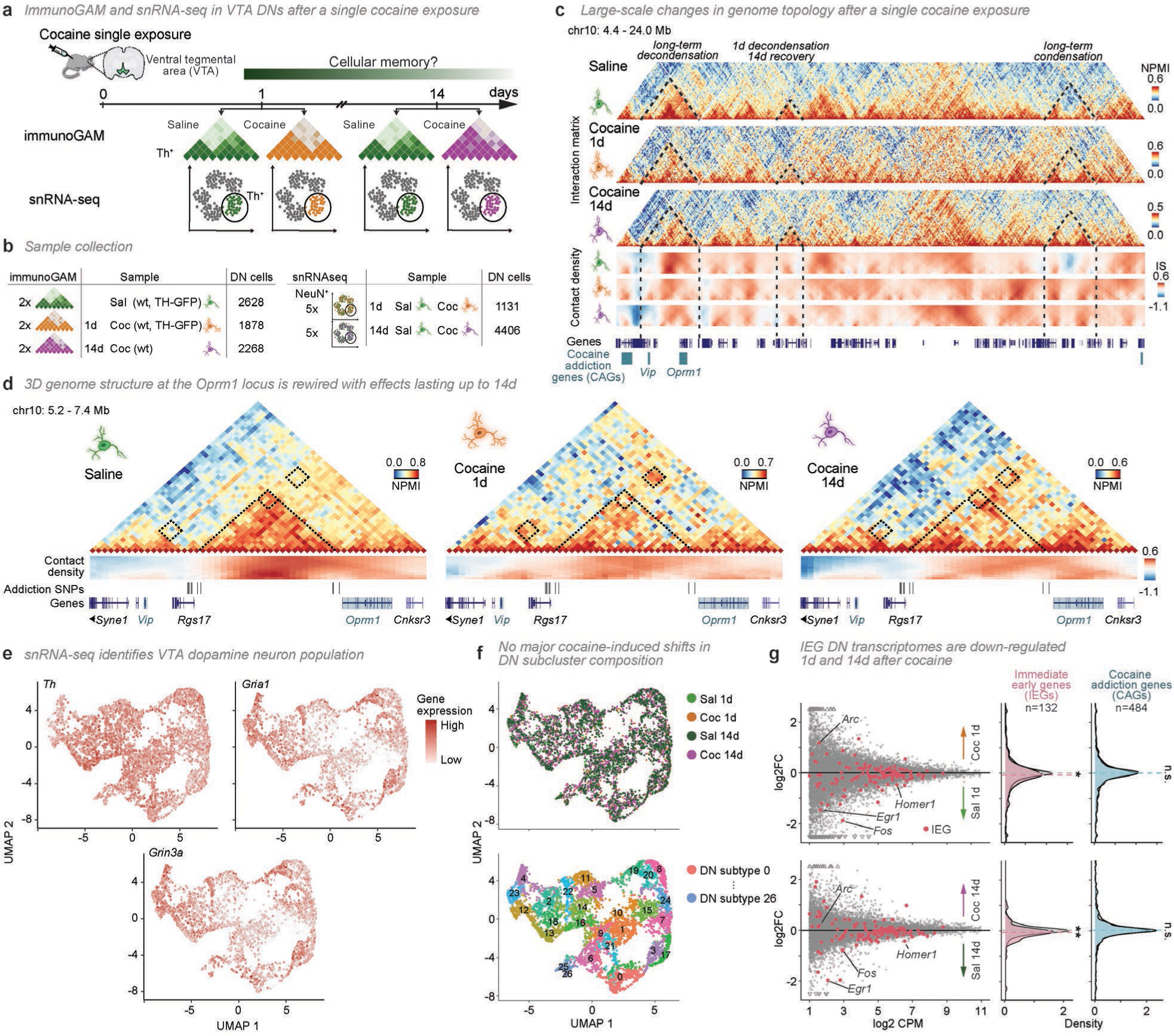
A single cocaine exposure induces large-scale disruption of 3D genome structure. **a**, 3D genome structure (immunoGAM) and single transcriptomes (single-nucleus RNA sequencing) in mouse dopamine neurons (DNs) from the ventral tegmental area (VTA), 1 or 14 days following single cocaine or saline exposure. **b**, Number of replicates (animals) and cells profiled. **c**, Example of long-term cocaine-induced 3D genome reorganization (chr10: 4.4 – 24.0 Mb; 40 kb resolution). NPMI, normalized pointwise mutual information. Contact density heatmaps represent insulation scores calculated with square boxes ranging 240-1040 kb (bottom to top, respectively). **d**, Cocaine-induced disruption of a domain demarcated by putative addiction single-nucleotide polymorphisms (SNPs) leads to long-term rewiring of flanking addiction-associated genes (chr10: 5.2 – 7.4 Mb). **e,** Representative Uniform Manifold Approximation and Projection (UMAP) expression profiles of DNs for a marker gene (*Th*) and other neuronal genes (*Gria1* and *Grin3a*). **f,** Projected distribution of cells according to treatment and DN subtype. **g**, Fold change representing differential gene expression 1- or 14-days following cocaine. No significant transcriptional differences of individual genes are detected 1- or 14-days following cocaine exposure, compared to saline treatment. Immediate early genes (IEGs) are highlighted in pink. The expression of IEGs, considered as a group, is downregulated 1 and 14 days after cocaine (two-sided Wilcoxon signed-rank test, *P* = 1.6 × 10^−2^ and 4.8 × 10^−3^, respectively; n.s., not significant). CAGs, Cocaine addiction genes.

To map 3D genome structure while avoiding tissue dissociation, we applied Genome Architecture Mapping with immunoselection (immunoGAM) to DNs in VTA samples, using tyrosine hydroxylase immunodetection as a marker. We collected immuno-GAM data from 4 cocaine-treated and 2 saline-treated animals, the latter datasets previously reported in Winick-Ng *et al.* 2021^10^. The GAM samples were produced from a total of 6774 DNs (4146 cocaine-treated and 2628 saline-treated; **Fig. 1b**, see **Extended Data Fig. 1a-d** for quality control metrics).

To characterise gene expression, we profiled single nuclear transcriptomes in the same conditions and time points. We collected VTAs from 20 wildtype mouse littermates, 1 or 14 days following treatment with saline or cocaine (**Fig. 1b**), and applied single nucleus RNA-seq (snRNA-seq)^17^ to neurons positive for the neuronal marker NeuN (**Extended Data Fig. 1e**), collecting high quality transcriptomes of 115,211 nuclei (see **Extended Data Fig. 1f-i** for quality control metrics)^18^.

### A single exposure to cocaine induces large-scale changes in DN chromatin topology

Inspection of chromatin contact matrices revealed many events of striking reorganization of 3D genome structure, at multiple genomic scales, upon cocaine exposure. For example, a representative 20 Mb region on chromosome 10 displays marked differences in chromatin contacts both at 1 and 14 days following cocaine exposure compared with saline injection (**Fig. 1c**, see **Extended Data Fig. 2a, b** for wildtype replicates and TH-GFP genotype). To assess whether genes previously associated with chronic cocaine exposure were amongst those affected by the extensive topological changes observed, we compiled a list of cocaine-associated genes (CAGs) from publicly available resources (**Supplementary Table 1**)^19,20^. At 1-day post-exposure, contact matrices show extensive losses and gains of contacts at both short and long genomic ranges with clear disruption of self-interacting topologically associating domains (TADs). By 14 days, remarkably, long-lasting differences in chromatin topology are evident, including at regions overlapping CAGs, such as *Vip*, involved in both neurotransmission and neuromodulation, and *Oprm1*, an opioid receptor gene (**Fig. 1c**). Some regions revert to their pre-cocaine state, while others show new structural alterations not present at 1 day.

To assess differences in contact density at different local scales, we measured insulation scores (average contact density) at genomic distances ranging 240 to 1040 kb, in saline, 1- and 14-day matrices^10,21^ (**Fig. 1c**, lower contact density panels). In some genomic regions, extensive decondensation seen 1-day post exposure reverts to pre-cocaine states by 14 days, while in many other regions, chromatin decondensation or condensation lasts or becomes more prominent by 14 days.

One exemplar locus with strong cocaine-induced reorganization contains the addiction genes *Oprm1* and *Vip*, separated by a large domain which is flanked by multiple putative conserved single nucleotide polymorphisms (SNPs) associated with cocaine and other comorbid addictions (**Fig. 1d**, see **Extended Data Fig. 2c, d** for replicates)^22,23^. The SNP list was compiled from publicly available resources and curated for human-mouse conservation (**Supplementary Table 1**)^19,24^. Cocaine exposure results in increased contacts between *Rgs17*, a modulator of the G-protein coupled receptor signaling pathway^25^, and *Cnksr3* or *Vip* after 1 day, which recovers after 14 days. In contrast, contacts between *Rgs17* and *Oprm1* are disrupted after 1 day, and do not recover even by 14 days. These observations show that a single exposure to cocaine results in changes in 3D genome topology that outlast transcriptional and synaptic effects, which occur throughout the genome including at genomic regions that contain addiction genes and non-coding addiction-associated SNPs.

To characterize gene expression genome-wide in VTA DNs treated in the same conditions, we selected a total of 5,537 DNs selected from VTA snRNA-seq transcriptomes using the marker genes *Th*, *Slc6a3, Slc18a2, Lmx1b, Foxa2, Nr4a2, Snca* and *Kcnj6* (**Extended Data Fig. 3a-i**)^26^. We confirmed DN identity by uniform expression of *Th*, which encodes tyrosine hydroxylase (**Fig. 1e**). Genes with synaptic plasticity-related functions, which are involved in cocaine-induced LTP^1^, were also highly expressed, such as *Gria1*, encoding an AMPA receptor subunit, or the NMDA receptor subunit *Grin3a* (**Fig. 1e**), while low expression of the cocaine response gene *Cartpt*^12^ indicated the DNs are not undergoing an active cocaine response (**Extended Data Fig. 3j**). We also find that the overall DN cluster substructure is not affected by cocaine exposure, and can be classified into 27 sub-populations, similar to recent DN subtype classifications^27^ (**Fig. 1f, Extended Data Fig. 3k, Supplementary Table 2**). We find that no individual gene is significantly differentially expressed 1 or 14 days following cocaine exposure, compared to saline treatment (**Supplementary Table 3**), in line with previous transcriptomic analyses performed 1 day after *in vivo* exposure to cocaine^9^. To search for more subtle long-term transcriptional changes, we considered the group of immediate early genes (IEGs), known to be upregulated within the first hours following neuronal activation after cocaine exposure^9^. We find a tendency for downregulation of IEGs at both 1 and 14 days, such as *Fos*, *Nr4a1* and *Homer1*, whereas CAGs, NMDA receptor (NMDAR) and AMPA receptor (AMPAR) genes are not affected (**Fig. 1g, Extended Data Fig. 3l-o**). These results suggest that a single cocaine exposure may result in a homeostatic response that overcompensates for the initial strong, cocaine-induced activation^28^.

### Cocaine exposure induces new TAD boundaries at postsynaptic genes

To investigate the extent of cocaine-induced large-scale changes in 3D genome structure and the affected genes, we started by comparing the organization of topologically associating domains (TAD)^10,21,29^. Cocaine exposure led to extensive genome-wide reorganization of TADs, with only 27% remaining unaffected by cocaine exposure, 49% appearing *de-novo* after cocaine exposure, 14% lost at 1 day but recovered at 14 days, and finally 10% lost and not recovered by 14 days (**Fig. 2a**). Remarkably, cocaine-induced TAD boundaries overlap with many neuronal genes, including IEGs (e.g. *Homer1*) as well as cocaine addiction genes (e.g. *Grin2b*, *Actb, Nlgn1*)^30–32^ (**Supplementary Table 4**), suggesting that the immediate transcriptional response to cocaine exposure leads to long-term domain reorganization that affects IEG responsive genes and cocaine addiction genes. We find that boundaries uniquely identified 1 or 14 days after cocaine exposure appear at neuronal-relevant genes, including the IEG *Bdnf*, or CAGs such as *Epha4*, *Grip1* and *Nlgn1*, and are characterized by gene ontologies (GOs) such as ‘synapse organization’ (e.g. *Lrk2* or *Bdnf*, respectively), ‘cell adhesion’ (at 1 day; e.g. *Ache*) or ‘localization within membrane’ (at 14 days; e.g. *Grip1*; **Fig. 2b, Extended Data Fig. 4a**).

**Figure 2.**
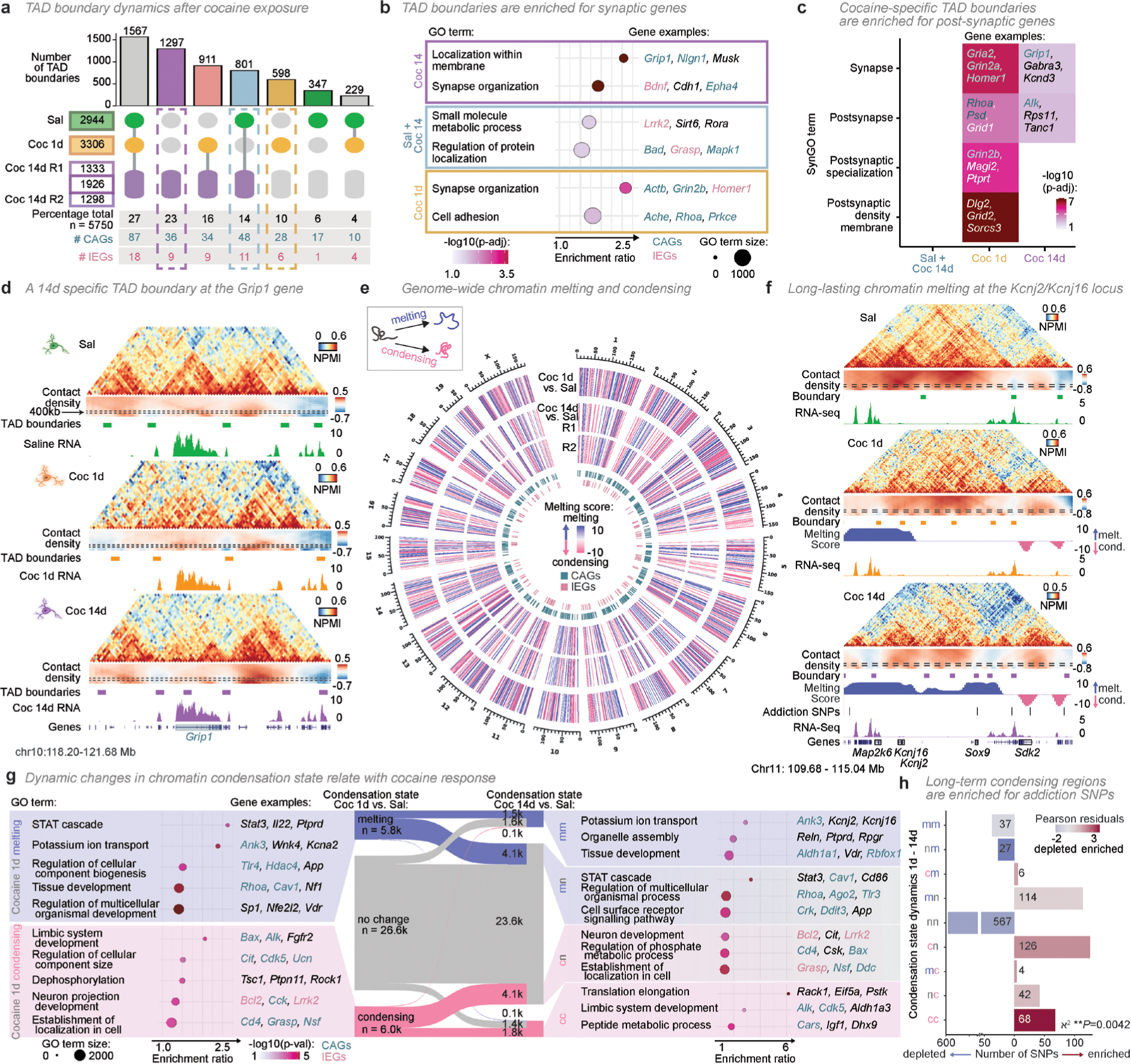
TAD borders and contact density are extensively rewired following a single cocaine exposure. **a**, UpSet plot of multi-way TAD boundary comparisons, considering 14-day boundaries found in either biological replicate. Sal, saline; Coc 1d, 1 day after cocaine; Coc 14d, 14 days after cocaine. **b**, Cocaine-response genes overlap cocaine-specific TAD boundaries. **c,** Genes overlapping cocaine-induced TAD boundaries have postsynaptic functions (synaptic gene ontology analysis; SynGO). **d**, *Grip1* overlaps a 14-day specific boundary (coloured boxes below contact density heatmap; chr10: 118 – 122 Mb). Dashed boxes on the contact density heatmap represent 400kb insulation scores, used to determine boundaries. Replicate 1 is shown for 14 days. **e**, Genome-wide melting and condensing, computed across a 120 kb sliding window, based on melting scores >5 or <-5, respectively (one-sided Kolmogorov–Smirnov test, *P* < 1 × 10^-5^). **f**, Example region showing melting of *Kcnj16* and *Kcnj2*, and condensing downstream of *Sox9*, at 1- and 14-days post-cocaine (chr11: 110 - 115 Mb). **g**, Melting and condensing dynamics following cocaine exposure, considering only events common to both 14-day replicates. **h**, Addiction-associated SNPs are enriched in condensing regions (*X*^2^ distribution test, \*\**P* = 0.0042). In **b** and **g**, top gene ontology (GO) terms were selected by adjusted *P*-value (p-adj) and enrichment ratio (observed over expected ratio of expressed genes).

Next, we applied SynGO (Synaptic Gene Ontology) enrichment analyses and found that TAD boundaries which are unique to cocaine exposure at 1 or 14 days were especially enriched for specialized postsynaptic functions (**Fig. 2c, Extended Data Fig. 4b**)^33^. Postsynaptic plasticity at glutamatergic synapses are largely responsible for the LTP effects observed after a single cocaine exposure^1,34^. For example, the TAD containing *Grip1,* a scaffolding protein gene critical for AMPAR trafficking during LTP^35^, is largely disrupted 1 day post-cocaine exposure, even though the flanking TAD borders remain stable (**Fig. 2d, Extended Data Fig. 4c, d**). By 14 days post-exposure, contacts are recovered only downstream of the first *Grip1* intron, establishing a smaller TAD and a new TAD boundary inside *Grip1*, while the first exon and intron remain highly decondensed. Together, these results show complex loss and gain of TAD boundaries, 1- and 14-days following cocaine exposure, which affect genes with synaptic functions and known roles in the cocaine-induced plasticity response, with potential long-term consequences for gene activity.

### Widespread changes in chromatin condensation upon cocaine exposure

Next, to quantify the broad changes in chromatin compaction genome-wide, we developed MELTRONIC (genomIC MELTRON), an approach based on the MELTRON pipeline which we previously developed to detect melting at long genes^10^. MELTRONIC quantifies gain (condensation) or loss (melting) of contacts genome-wide by applying a sliding window of differential insulation scores across the genome. Using insulation boxes from 240 to 1040 kb, and a sliding window of 120kb, MELTRONIC detected chromatin melting and condensation on all chromosomes at both 1 and 14 days in comparison with saline (**Fig. 2e, Extended Data Fig. 5a, b, Supplementary Table 5**). Condensing and melting regions were defined using conservative melting score thresholds of −5/5 (equivalent to an adjusted *P* < 1 × 10^−5^), in line with previous work^10^ (**Extended Data Fig. 5c, d**). Melting and condensing events are stronger at 1 compared to 14 days after cocaine exposure, but many genomic regions remain or become *de-novo* melted or decondensed by 14 days. The prevalence of melting and condensing states at 14-days post exposure are confirmed by separate analyses of the two biological replicates, which showed high conservation (61.6%, **Extended Data Fig. 5e**).

Genomic regions affected by long-term changes in chromatin compaction after cocaine exposure include the genomic regions containing the *Kcnj2* and *Kcnj16* locus, two potassium ion channel genes that modulate LTP sensitivity^36^, as well as several addiction-associated SNPs (**Fig. 2f, Extended Data Fig. 5f, g**). The region also contains *Sox9*, a neural stem cell fate gene, which has been extensively studied in the context of genetic rearrangements that alter 3D genome topology and gene expression, leading to human developmental diseases^37,38^. We show that a single exposure to cocaine results in the loss of the TAD boundary separating *Kcnj2/16* from *Sox9* and its enhancers by 1 day post exposure, coinciding with melting events upstream of *Kcnj2/16*, across a large 1.5 Mb genomic region that also contains *Map2k6*, a mitogen-activated protein kinase gene, and other highly expressed genes. By 14 days, chromatin melting extends downstream of *Kcnj2/16*, across the region surrounding the *Sox9* locus. Long-term condensation events are also observed in the same region and affect for example *Sdk2*, a paralog of *Sdk1* which is upregulated after chronic cocaine use in the Nucleus accumbens (NAc)^39^. These long-lasting chromatin compaction changes suggest the long-term cocaine-induced propagation of melting and condensation of genes related to neuronal activity, including at many SNP containing regions.

Next, we performed GO analyses to explore which genes are affected by the different dynamics of melting/condensing at 1- and 14-days post cocaine exposure (**Fig. 2g**). For example, 1-day melted regions contain genes with functions in the STAT cascade and potassium ion transport, with the latter typically remaining melted at 14 days. Condensing regions at 1-day post exposure are enriched for genes associated with neuron projection development and limbic system development, with the latter remaining condensed at 14 days. SynGO enrichment analyses identify genes with roles in synaptic organisation enriched in 1-day melting regions and in 14-day condensing regions (**Extended Data Fig. 5h**). For example, the receptor tyrosine kinase *Alk*, which concentrates in post-synaptic domains and contains multiple addiction-associated SNPs in its intronic regions^40^, condenses at 1 day and remains condensed at 14 days (**Extended Data Fig. 5i, j**). Interestingly, 1- and 14-days condensing regions are also significantly enriched in addiction-associated SNPs (*P* = 0.0042, Chi-squared test; **Fig. 2h**). Taken together, long-term chromatin melting and condensing are widespread cocaine-associated phenomena that affect a significant fraction of the genome, occurring both at coding regions of genes with important transcriptional and synaptic functions, and at non-coding regions with regulatory or structural roles.

### Long neuronal genes undergo cocaine-induced melting and condensing events

Long genes are more highly expressed in terminally differentiated cells, including neurons, where their melting state reflects their expression levels in specific neuronal cell-types^10,42–44^. To quantify the effects of cocaine exposure on the folding of long genes, here longer than 280 kb, we applied MELTRON^10^ and found that approximately half undergo melting or condensing events at 1 day post-cocaine exposure (140 out of 291 genes; **Fig. 3a, Supplementary Table 5**). Amongst these genes, 73 retain their melted/condensed state at 14 days, while an additional 46 genes melt or condense *de novo* (19 and 27 genes, respectively). Of interest, many of the genes that undergo cocaine-induced melting in DNs were previously found melted in pyramidal glutamatergic neurons (PGNs; e.g. *Rbfox1*)^10^, suggesting that cocaine alters the cell-type specificity of chromatin condensation.

**Figure 3.**
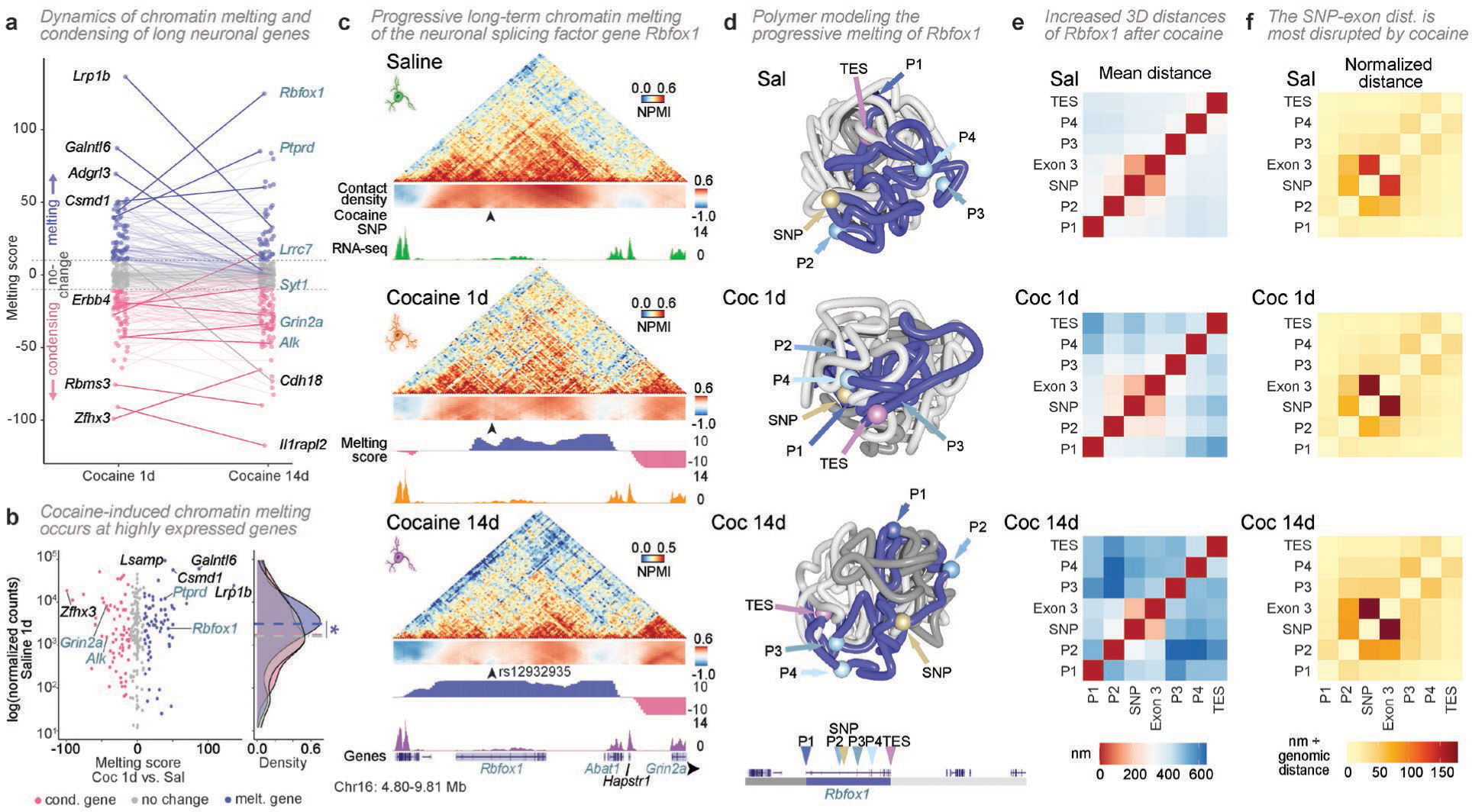
*Rbfox1* melting is more pronounced 14 days after cocaine exposure. **a,** Melting scores from long (> 280kb) expressed genes, 1 or 14 days following cocaine. **b**, Melting at 1 day of cocaine is associated with higher baseline transcription (two-sided Wilcoxon signed-rank test, \**P* = 0.02). Expression is shown for the matching samples collected 1-day after injection with saline. **c**, Example region showing melting of *Rbfox1*, 1 and 14 days after cocaine (chr16: 4.8 - 9.8 Mb). Arrowhead indicates a putative cocaine SNP. **d**, Polymer models of the *Rbfox1* region show looping near the transcription end site (TES) at 1 day, and full gene decondensation by 14 days. Colour bars, gene region, up- and downstream flanking regions. Coloured spheres and arrowheads, positions of promoters and SNP. Cocaine-induced increased 3D spatial separation of promoters and SNP **e**, especially between the SNP and downstream exon after normalization of linear genomic distances **f**.

Long genes that are melted 1-day post-cocaine exposure tend to be more highly transcribed in saline-treated cells compared to genes that condensed or are unchanged, suggesting that higher transcription rates sensitize genes to larger scale melting events (**Fig. 3b**). By 14 days post exposure, the melting status is much less related with gene expression in saline treated cells (**Extended Data Fig. 6a**) and is possibly a bystander consequence of the initial direct effects of cocaine-induced transcriptional activation on chromatin topology.

### The *Rbfox1* gene melts extensively 14 days after cocaine exposure

To explore further the long-term effects of cocaine exposure on chromatin folding, we focused on *Rbfox1*, a cocaine addiction gene which regulates alternative splicing following multiple cocaine treatments in the NAc^45^, and is the most melted gene at 14 days post cocaine exposure. In saline-treated DNs, *Rbfox1* is contained within two large chromatin domains with many inter-TAD interactions (**Fig. 3c**, **Extended Data Fig. 6b** for a comparison of the two 14-day replicates). At 1 day after cocaine exposure, *Rbfox1* is extensively decondensed (melting score = 43), especially in its 3’ end. By 14 days, its melting score increases further (to 125), especially at an intronic region containing a putative cocaine-associated SNP (rs12932935). The SNP coincides with putative transcription factor (TF) binding sites for IEGs, including *Jun* and *Stat1*, as well as the circadian factor *Nr1d2* (**Extended Data Fig. 6c**), suggesting that LTP-induced TF activity may be targeted at, or involved in, the long-term structural reorganization of chromatin after a single cocaine exposure.

To further understand the cocaine-induced structural changes in the *Rbfox1* locus, we generated ensembles of 3D models (1100 models per condition), using a polymer-physics based approach^10,46^ which were validated by reconstructing *in-silico* GAM matrices (**Extended Data Fig. 6d**). Inspection of single models shows that different sections of the *Rbfox1* gene are more tightly compacted in saline, with proximal positioning of several *Rbfox1* promoters and its TES (**Fig. 3d**; see more examples in **Extended Data Fig. 6e, Supplementary Videos 1-6**). At 1-day post-exposure, the genomic region between the promoter 4 and TES of *Rbfox1* loops out from the rest of the polymer (**Supplementary Videos 3 and 4**). By 14 days, the entire *Rbfox1* gene becomes highly extended, with a larger separation between all its promoters, the cocaine SNP, and the TES. The cocaine-induced increased separation between promoters and SNP, especially at 14 days, are confirmed across the ensemble of polymers (**Fig. 3e, Extended Data Fig. 6f, g**). Variance analyses showed that these internal distances are most highly variable across polymers after 14 days (**Extended Data Fig. 6g**), indicating that the single exposure to cocaine results in loss of structural coherence in the 3D organization of *Rbfox1*. After normalizing for linear distances, we found that the most pronounced increases in physical distance upon cocaine exposure are between the SNP, its closest exon, the upstream internal promoter P2 and to a smaller extent P3 (**Fig. 3f**). This suggests that the post-cocaine structure of *Rbfox1* may sensitize it to future activation, in particular at some of its internal promoters, and highlights a potential role of genetic variation in the SNP neighbourhood in the magnitude of topological changes occurring with a single cocaine exposure.

We were surprised by the similarity of structural properties of the *Rbfox1* locus found at 14 days post-cocaine exposure with that in hippocampal pyramidal neurons (**Extended Data Fig. 6h**), a cell type with much higher baseline expression of *Rbfox1* than DNs^10^. When modeled in the pyramidal neurons, *Rbfox1* polymers have distance and variation similar to the cocaine-treated DNs at 14 days (**Extended Data Fig. 6h, i**), suggesting that *Rbfox1* was either previously activated in DNs by the cocaine exposure and/or may become sensitized to future activations. Together, these data suggest that, by 14 days, the 3D genome structures of specific loci are not trivially recovering their pre-drug state following cocaine exposure, but can additionally undergo further structural rewiring, including loss of structural coherence, consistent with a progressive cascade of disruption.

### Compartment A/B transitions affect one third of the genome and include CAGs

To understand whether melting dynamics relate to broader scales of 3D genome organization, we calculated compartment A/B classifications, reflecting open and closed chromatin, respectively^10,41^. Compartment changes occurred in 29% of the genome and also highlighted long-term effects of a single cocaine exposure (**Extended Data Fig. 7a, b**). For example, compartment transitions from A-B-A (saline to 1 day to 14 days) are enriched for signaling genes (e.g. *Snca*, *Sst*), A-A-B transitions for genes with roles in cell adhesion and response to external stimulus (e.g. *Clock*, *Tyr*), and A-B-B transitions for synaptic transmission genes (e.g. *Oprm1*, *Gabra2*), many of which are CAGs (**Extended Data Fig. 7c**). As shown previously^10^, compartment A/B changes are mostly independent of melting and condensing events (**Extended Data Fig. 7d**). These results show that compartment transitions are found at relevant regions for the cocaine response, though they occur less frequently than TAD reorganization and are independent of (de)condensation events.

### Strong cocaine-specific contact regions involve cocaine-response genes and IEGs

Finally, we also explored more complex structural changes across genomic regions spanning several megabases, such as those seen at the clustered proto-cadherin locus encoding cell adhesion genes (*Pcdh α*, β, and γ)^47^, which show increased contacts 1 day post-exposure with gene-dense regions up- and downstream that include CAGs (**Fig. 4a**). To unbiasedly discover other ‘hotspot’ genomic windows characterized by an excess of differential contact loops, we calculated differential matrices between cocaine and saline treatments and determined the number of differential loops that each genomic region establishes within 2.5 Mb distance, using a previously reported approach^10,29^ (**Supplementary Table 6**). The whole *Pcdh* cluster is detected as a hotspot of differential contacts 1-day post exposure, especially the β cluster which also remains a hotspot of cocaine-specific contacts at 14 days post-exposure (**Fig. 4b**).

**Fig. 4.**
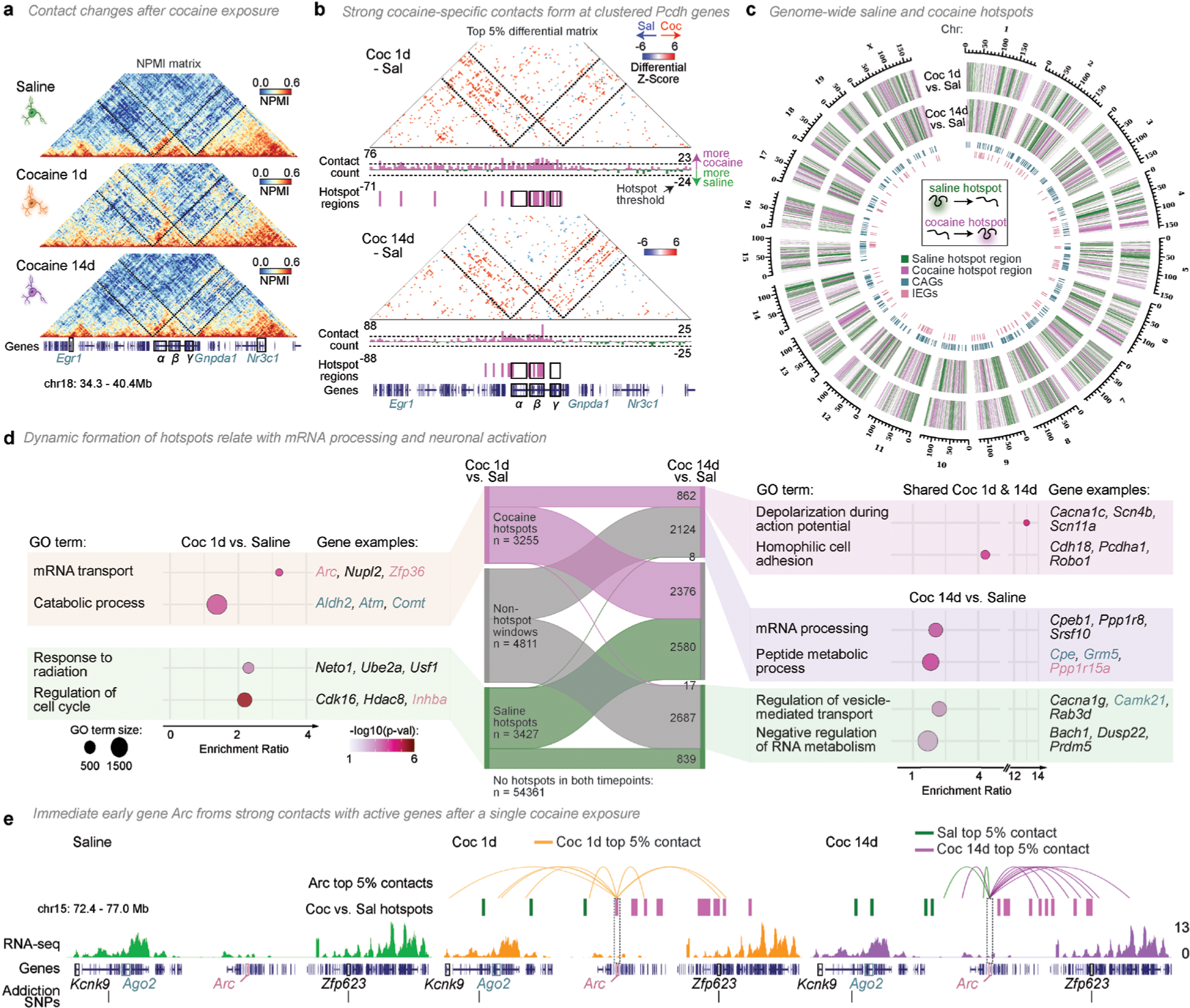
Hotspots of strong structural changes occur at the *Pcdh* cluster, mRNA processing genes and *Arc*. **a**, Strong cocaine-specific contacts are formed in windows containing the clustered protocadherin genes (*α*, β, and γ clusters; chr18: 34 - 40 Mb). **b**, Hotspot regions, top 5% of summed genomic windows containing differential contacts (dashed lines below matrix). **c**, Genome wide hotspots. **d,** Hotspot dynamics following cocaine exposure, and related GO enrichments. **e**, Example hotspot at *Arc* 1 day following cocaine (chr15: 72 - 77 Mb). *Arc*-anchored, treatment-specific, contacts are shown with orange (1 day cocaine), purple (14 day cocaine) and green (saline) lines below the hotspot track.

When extended genome-wide, the contact hotspot analysis shows that the genomic regions most affected by cocaine treatment are often clustered along the linear genome (**Fig. 4c, Extended Data Fig. 8a**). Chromosome 18, for example, has long contiguous stretches of hotspot windows within a given treatment (**Extended Data Fig. 8b**). The average genomic length of contiguous hotspots of differential contacts ranges from 1.9 - 2.6 Mb (± 0.1 - 0.2 mean standard error) and increase in length after 14 days (**Extended Data Fig. 8c**), suggesting the local propagation of chromatin structure disruption between 1- and 14-days post exposure. Many cocaine-specific hotspots are maintained between 1 and 14 days (n=862/3254) and are enriched for genes related to membrane depolarization and cell adhesion, including the clustered *Pcdh* genes (**Fig. 4d**). Interestingly, cocaine-associated hotspots both at both 1 and 14 days also include genes involved in mRNA transport, splicing and processing, consistent with widespread alternative splicing of pre-mRNAs described in VTA and NAc following chronic self-administration of cocaine^48^.

Cocaine-associated hotspots also include several IEGs, such as the memory- and stress-associated genes *Arc, Hspa4*, *Ppp1r15a*, and *Zfp36* (**Extended Data Fig. 8d**)^49–52^. The 1-day cocaine-specific hotspot at *Arc* is of particular interest, as *Arc* is directly linked to the reinforcing properties of cocaine exposure and has been referred to as the ‘master organizer of long-term synaptic plasticity^53^. We observed that *Arc* gains contacts with upstream active genes specifically at 1-day post cocaine exposure, including the addiction-associated gene *Ago2*^54^, 1.5-Mb away, and the zinc-finger transcription factor *Zfp623*, 1.3-Mb away, which contains a putative alcohol-dependence SNP^55^ (**Fig. 4e**). Though after 14 days, *Arc* is no longer within a hotspot region, it maintains its strong contacts with *Zfp623*. These results suggest that IEGs, a group of genes that remained down-regulated 14 days after a single cocaine exposure (**Fig. 1g**), are present at genomic regions that acquire extensive cocaine-specific contacts, including specific long-lasting contacts with critical cocaine- and addiction-associated risk genes.

### A DN sub-cluster defined by IEGs and genes with chromatin contact changes localizes to the medial VTA

Having found extensive 3D genome structural changes at IEGs, CAGs and many other genes upon cocaine exposure, we explored whether these genes were expressed in specific DN subtypes, especially because previous work has shown that DNs in different VTA subregions respond to different extents to a single administration of cocaine *in vivo*^56^, and project to secondary addiction regions^57–59^. We started by asking whether IEG expression, and their downregulation by 14 days after cocaine exposure, was common across the population of DNs, or specific to a DN subtype, by plotting their expression on the DN UMAPs. We found that some IEGs, such as *Arc*, *Egr1*, *Fos*, and *Homer1*, are specifically and/or more highly expressed within a small DN sub-type in saline conditions (212/5537 DNs; **Fig. 5a**). This DN sub-population shows a high combined expression of IEGs, and when compared with all other DN subtypes, it is largely responsible for the observed down-regulation of IEGs after cocaine exposure (**Fig. 5b**; see full DN UMAP in **Extended Data Fig. 9a**).

**Figure 5.**
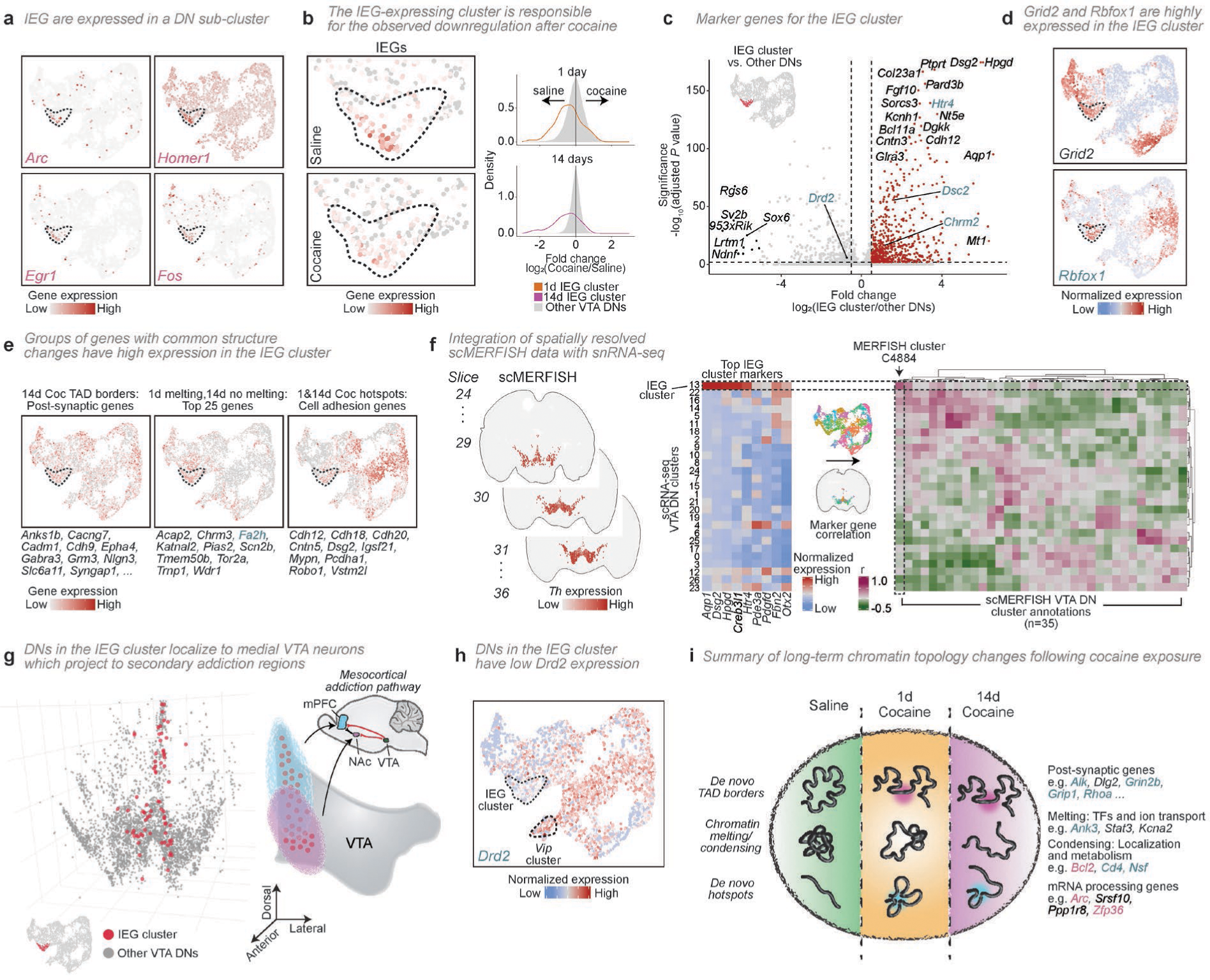
A DN sub-cluster localizes to the medial VTA, expresses genes with chromatin contact changes, and displays long-term IEG downregulation. **a,** UMAPs of example IEG expression in DNs. Dashed line indicates a DN sub-cluster with higher expression of indicated genes. **b**, IEGs are highly expressed in the DN sub-cluster, but downregulated after cocaine (Permutation test, P = 1.3 × 10^−9^ and 1.2 × 10^−19^ for 1d and 14d, respectively). Density plots show IEG expression in the ‘IEG cluster’ compared to all other DNs. **c**, Volcano plot of differentially expressed genes in the IEG cluster. Red dots indicate marker genes with higher expression in the cluster (Wilcoxon test, *P* < 0.05, fold-change > 1). UMAPs of **d**, individual examples or **e**, groups of genes with cocaine-induced chromatin structural changes that have high expression in the IEG cluster. **f**, Integration of sn-RNA-seq with single-cell MERFISH (scMERFISH)^27^ and identification of the IEG cluster in scMERFISH by correlating top IEG cluster marker genes to scMERFISH cluster annotations. **g**, IEG cluster DNs (pink dots) localize to the medial VTA. The medial VTA projects to the nucleus accumbens (NAc) and prefrontal cortex (mPFC)^60,61^. **h,** UMAP of *Drd2* expression showing low expression in the IEG cluster and high expression in the *Vip* cluster (dashed lines correspond to cluster annotations on the left). Low *Drd2* expression in midline VTA DNs is associated with increased LTP sensitivity after cocaine exposure^56,61^. **i**, Summary of long-term 3D genome structural changes after a single cocaine exposure.

To characterize the ‘IEG-expressing’ cluster of DNs, we inspected their marker genes and found many associated with addiction, such as *Chrm2*, *Dsc2*, *Hpgd*, *Htr4*, and *Nt5e* (**Fig. 5c, Extended Data Fig. 9b**; see cluster 13 in **Supplementary Table 2**)^57–59,62,63^. Importantly, the IEG cluster of DNs also shows high expression of many addiction-associated genes which we found to undergo cocaine-induced chromatin rewiring. Some examples include *Rbfox1*, with its extensive melting 14 days after cocaine; *Grid2*, with very strong hotspots of differential contacts in saline which are lost at both 1 and 14 days; and *Ptprt*, which condenses at 1 and 14 days (**Fig. 5d, Extended Data Fig. 9c-e**). Other groups of genes with chromatin structural changes also showed higher expression in the IEG cluster, including genes with the strongest melting scores 1 day after cocaine, cell adhesion genes found within 1- and 14-day cocaine hotspots, and synapse organization and postsynaptic genes found at new cocaine TAD borders (**Fig. 5e**, see **Extended Data Fig. 9f** for other example groups).

### The IEG-expressing DNs locate to the medial VTA and have features of highly cocaine-sensitive cells

To investigate the VTA localization of the IEG-expressing DNs which more highly express genes that undergo chromatin structural changes after cocaine exposure, we took advantage of a recent MERFISH single-cell spatial transcriptomic atlas in the mouse brain, based on the expression of 500 genes^27^. We identified the brain slices containing the VTA based on MERFISH annotations, and then correlated the expression of DN marker genes defined in the present study with the VTA MERFISH clusters (**Fig. 5f, Extended Data Fig. 10a**). The IEG-expressing DNs could be robustly matched to a specific MERFISH cluster corresponding to 2% of all VTA DNs (96/4115 DNs; **Fig. 5f**). Remarkably, we found that the IEG-expressing cluster of DNs specifically localize along the entire VTA midline (**Fig. 5g, Supplementary Video 7**). Previous work showed that DNs located in the ventral and dorsal midline of the VTA project to the NAc and medial prefrontal cortex, respectively ^60,61,64^, which are two regions associated with cocaine craving and relapse^65,66^. As a control, we confirmed that the cluster of *Vip*-expressing DNs could also be correctly assigned to the dorsal midline, as expected^67^ (**Extended Data Fig. 10b-d, Supplementary Video 8**). Midline DNs which have the strongest cocaine response, characterized by high postsynaptic LTP sensitivity, are known to project to the medial shell of the NAc and are characterized by low expression of the dopamine auto-receptor *Drd2*. We found that *Drd2* is lowly expressed in the IEG cluster of DNs compared with all other VTA DNs, and more highly expressed in the *Vip* cluster (**Fig. 5c, h**), suggesting that the small IEG-expressing cluster of DNs may be the most sensitive population of VTA DNs to a single cocaine exposure.

## Discussion

Upon a first drug exposure, neurons undergo a strong, though transient, transcriptional and physiological response; however, where the cellular memory of that exposure is stored is unknown. In this study, we discovered that a single dose of cocaine is sufficient to induce large-scale reorganization of chromatin structure that lasts far past the initial physiological response. We showed that genome rewiring occurs across a broad spectrum of genomic distances and regions, with extensive chromatin structural changes that affect numerous genes, including many associated with cocaine, addiction, or synaptic plasticity (**Fig. 5i**). Chromatin reorganization seen after 1 day of cocaine exposure often persists or can appear *de novo*, 2 weeks later, well past reported transcriptional changes^4–7^ and LTP effects^9^, highlighting an unexpected long-lasting storage of the effects from a first exposure to highly addictive drugs in chromatin structures.

Our findings support the involvement of gross changes in 3D genome structure and the hypothesis of long-term ‘chromatin memory’ storage in the inception of drug addiction. *In vitro* studies have shown that chromatin sites can remain locally open either by TF retention or new TF binding following synchronous neuronal activation, though gene expression returns to its pre-activated state by 24 hours^13,68^. Expanding from the long-lasting TF-related chromatin regulation, we find extensive reorganization of the 3D structure of genomic regions containing genes that encode for neuronal activation TFs at 14 days post cocaine exposure, such as *Stat* genes in melting regions. We also find traces of long-lasting changes related with IEG activity, for example the presence of putative *Jun* binding sites at *Rbfox1* internal promoters and intronic SNP, which are extensively decondensed 14 days post single exposure. The widespread chromatin reorganization, across genomic scales, induced by cocaine exposure suggests a larger contribution of chromosome structure to the cellular memory of drug exposure than previously expected. Other cell-extrinsic mechanisms are also likely to contribute to the cellular memory and long-term chromatin topology changes, including nuclear involutions so far reported following *in-vitro* cocaine exposure^69^, homeostatic-related nuclear volume changes seen in epidermal self-renewal^70^, or changes to the physical state of chromatin due to altered nuclear ion concentrations^71,72^. Future work will be critical to tease apart both the cell-intrinsic and cell-extrinsic mechanisms which lead to long-term chromatin changes induced by a single exposure to cocaine, as well as efforts to explore potential recovery mechanisms to pre-exposure states.

Midbrain DNs are a highly diverse cell type, which connect to different parts of the brain with major roles in chronic addiction, such as the NAc and medial prefrontal cortex^60,61,64^. Each DN subtype has low abundance, for example, 96 IEG-expressing DNs were reported in the MERFISH database, out of 4115 VTA DNs, which pose further challenges in future research to understand their role in the inception of drug addiction. The knowledge of the genes that are more extensively structurally remodelled, obtained through GAM analyses, provides new insights into the long-term effects of a single exposure to cocaine, for example by enabling the identification a specific sub-type of DNs that more highly express these genes, as well as IEGs, and the discovery that IEGs are downregulated at both 1 and 14 days after cocaine exposure, especially in the same group of DNs. Given the widespread cocaine-induced chromatin changes observed in GAM data collected from the full population of VTA DNs, it is unlikely that chromatin reorganization is restricted to the IEG sub-cluster. However, the genome structural changes within the IEG-expressing DNs may make them more sensitive to further cocaine-induced topological alterations upon repeated usage, especially in genome regions containing genes with important addiction-related roles, such as in the postsynaptic LTP response, which are at baseline more expressed in the IEG cluster than in other DNs. The IEG-expressing DNs weakly express the dopamine auto-receptor, a feature of DNs with a highly sensitized cocaine response, and localize to midline VTA areas, which are known to project to secondary addiction brain regions^56,61^. Further developments and targeted experimental designs will be required to specifically study the very small populations of IEG-expressing DNs towards understanding whether they undergo more severe cocaine-induced chromatin changes compared with DNs that have different cocaine responses (i.e. inhibition following reward learning)^73^, and to further explore how cocaine-induced changes in 3D genome structure may specifically sensitize DNs that are highly activated by cocaine.

There are some important limitations to the present study. The analysis of two different genotypes in saline and cocaine, 1-day post-exposure, had value to show that cocaine-induced disruption of chromatin structure occurs in both genotypes, which also revealed locus-specific effects. Further work is needed to more broadly explore the effects of genotype, individual variation, age and sex, as well as the cascade of changes in 3D genome structure occurring before 1 day, until 14 days, and beyond. We profiled ∼5,500 DN transcriptomes from 20 animals, but their unexpected and extensive subtype variability inevitably results in low cell representation in each subcluster which limits the discovery of subcluster-specific changes in gene expression upon cocaine. Our study supports the need for future large-scale projects to dissect the effects and mechanisms of cocaine exposure on DN subtypes and other VTA cell types, as well as other brain regions containing DNs.

Our work opens many new questions, including: Can chromatin structure eventually recover to the pre-cocaine state, and is there a ‘critical wait period’ that might avoid cumulative effects of subsequent exposures? Importantly, can the wait period be shortened by external interventions? How does long-term chromatin remodelling after a single cocaine exposure contribute to drug seeking or susceptibility to addiction? Does the genetic and 3D genome make-up of individuals impact their likelihood for a stronger disruption of chromatin structure and of developing addiction? More broadly, our work also suggests the need to study the effects of other addictive drugs on 3D chromatin structure, including drugs which target the reward-learning (e.g. nicotine, amphetamine, alcohol) or other (e.g. heroin or fentanyl) pathways.

Our work identifies unexpected long-term effects of drug usage on 3D chromatin structure across the genome, including at drug-response genes, which may be crucial in the homeostatic responses of specific DNs. It highlights the plasticity of 3D genome structure in the context of addiction, as chromosomes are large physical objects with specific structure which, when disturbed, may require long periods of time to re-establish their healthy conformations, especially in non-dividing terminally differentiated cells such as neurons. By mapping 3D genome structure, we open new avenues to identify critical pathways and targets for intervention in the progression towards addiction.

## Acknowledgments

The authors thank Sheila Q. Xie and Azhaar Ashraf for help processing midbrain samples; members of the Pombo laboratory for helpful discussions and manuscript feedback; Thomas Conrad and Cornelius Fischer of the MDC/BIH Genomics Platform for sequencing, as well as scientific and technical support, and Anja Schütz and her team of the Protein Production and Characterization Platform, both at the Max Delbrück Center for Molecular Medicine in the Helmholtz Association, Berlin; Francesco Musella for guidance in using the in-silico GAM algorithm; Irina Korshunova of the BRIC single cell genomics core facility for assistance and support in the single-nucleus RNA-sequencing; Ben Blencowe for his careful feedback and editing of the manuscript; and Gonçalo Castelo-Branco for helpful discussions and advice.

## Author contributions

Conceptualization: A.P., M.A.U., W.W.-N.; Data Curation: D.S., W.W.-N., V.F., S.B., M.Y.B., A.K., S.D.; Formal Analysis: D.S., W.W.-N., V.F., S.B., M.Y.B, A.K., I.I-A., A.M.C, J.P.L.-A.; Funding Acquisition: A.P., M.A.U., A.A., K.K., M.N.; Investigation: W.W.N., M.Y.B., E.J.P, U.P., A.P.; Methodology: W.W.-N., D.S., V.F., A.P., S.B., A.A., K.K., M.N., U.P.; Project Administration: W.W.-N., A.P., D.S.; Resources: A.P., A.A., K.K., M.N., M.A.U.; Software: D.S., V.F., W.W.N, S.B., A.K., I.I-A, A.M.C., M.Y.B, S.D.; Supervision: A.P., W.W.-N, A.A., M.A.U, K.K., M.N.; Validation: V.F., D.S., W.W.-N., A.P., L.Z.-R., S.B., M.Y.B., E.J.P, A.K., I.I-A, A.M.C, S.D., K.K., M.A.U.; Visualization: W.W.-N., D.S., A.P., V.F., S.B., M.Y.B., A.K.; Writing – original draft: W.W.-N., D.S., A.P.; Writing – review & editing: A.P., W.W.-N., D.S., V.F., M.A.U., E.J.P., L.Z.-R., M.Y.B, S.B., A.K., J.P.L-A.

## Materials & Correspondence

Correspondence and requests for materials should be addressed to.

## Methods

### Animal maintenance

GAM and snRNA-seq data was collected using C57Bl/6NCrl (RRID: IMSR_CR:027; WT, Charles River), and TH-GFP (B6.Cg-Tg(TH-GFP)21-31/C57B6) male mice as previously described^10^. All procedures were approved by the Imperial College London’s Animal Welfare and Ethical Review Body, and were conducted in accordance with the Animals (Scientific Procedures) Act of 1986 (UK). All mice had access to food and water *ad libitum* and were kept on a 12 h:12 h day/night cycle in social groups of 3-4, with appropriate environmental enrichment. C57Bl/6NCrl and TH-GFP mice received an intraperitoneal (IP) injection of either saline or cocaine (15mg/kg body weight) 1 or 14 days or prior to the tissue collection. Cocaine-treated and saline-treated mice were housed in separate groups. Mice used for GAM and RNA-seq experiments were littermates. All mice were the same age at the time of collection (8 weeks old) and sacrificed in parallel.

### Tissue fixation and preparation for GAM

Tissue was prepared for GAM as previously described^10^. Briefly, mice were anesthetized with isoflurane (4 %), given a lethal IP injection of pentobarbital (0.08 μl; 100 mg/ml; Euthatal), and perfused with ice-cold phosphate buffered saline (PBS) followed by approximately 100 ml of 4% depolymerised paraformaldehyde (PFA; Electron microscopy grade, methanol free) in 250 mM HEPES-NaOH (pH 7.4-7.6; PFA-HEPES). Following perfusion, brains were removed and the VTA was isolated before quickly transferring to 4% PFA-HEPES for 1 h at 4°C, followed by 2-3 h in 8% PFA-HEPES. Tissue was placed in 1% PFA-HEPES at 4°C until prepared for cryopreservation.

### Cryoblock preparation and cryosectioning

VTA tissue samples were further dissected to produce ∼3×3 mm tissue blocks suitable for Tokuyasu cryosectioning^10^. Tissue blocks were post-fixed for 1 h at 4°C in 4% PFA-HEPES, before being transferred to 2.1 M sucrose in PBS for 16-24 h at 4°C. Sucrose-embedded blocks were mounted on copper stub holders before being flash frozen in liquid nitrogen. Tissue blocks were cryosectioned, as previously described^10^, with an Ultracryomicrotome (Leica Biosystems, EM UC7) at a thickness of 220-230nm and transferred onto a 4.0 µm polyethylene naphthalate (PEN; Leica Microsystems, 11600289) membrane for laser microdissection.

### Immunofluorescence detection for laser microdissection

Cryosections on PEN membranes were washed 3x in PBS, quenched with 20mM glycine in PBS for 20 min, then permeabilized with 0.1% Triton X-100 in PBS. After blocking for 1 h at room temperature in blocking solution (1% BSA (w/v), 0.2% fish-skin gelatin (w/v), 0.05% casein (w/v) and 0.05% Tween-20 (v/v) in PBS), cryosections were incubated in primary antibody overnight at 4°C with sheep anti-TH (1:50; Pel Freez Arkansas, P60101-0), followed by 3-5 washes for 1h in blocking solution at room temperature. Cryosections were incubated for 1h in secondary antibody (1:1000 donkey anti-sheep conjugated with AlexaFluor-488; ThermoFisher Scientific (Invitrogen)), followed by 2 washes in 0.5% Tween-20 in PBS and 1 wash in water. After drying, cryosections were visualized as previously described^10^, using a Leica laser microdissection microscope (Leica Microsystems, LMD7000) with a 63x dry objective. TH-positive nuclei from cellular sections (nuclear profiles; NPs) were laser microdissected, and collected into PCR adhesive caps (AdhesiveStrip 8C opaque; Carl Zeiss Microscopy #415190-9161-000). Three NPs were collected into each cap, with control lids not containing NPs (water controls) included for each dataset. In one experiment (1 day cocaine exposure in the TH-GFP mouse), 34 single NPs were collected into separate caps and later combined *in-silico* into 3 NPs (see ***GAM data window calling*** below; **Supplementary Table 7**).

### Whole genome amplification of nuclear profiles

Whole genome amplification (WGA) was performed as previously described^10^.

Briefly, NPs were lysed for 4 or 24 h at 60°C in lysis buffer (final concentration: 30 mM Tris-HCl pH 8.0, 2 mM EDTA pH 8.0, 800 mM Guanidinium-HCl, 5 % (v/v) Tween 20, 0.5 % (v/v) Triton X-100) and 2.116 units/ml QIAGEN protease (Qiagen, 19155), before a 30 min protease inactivation at 75°C. Pre-amplification was done using a 2x DeepVent mix (2x Thermo polymerase buffer (10x), 400 µM dNTPs, 4 mM MgSO_4_ in ultrapure water), 0.5 µM GAT-7N random hexamer primers with an adapter sequence and 2 units/µl DeepVent^®^ (exo-) DNA polymerase (New England Biolabs, M0259L). DNA was further amplified as in the pre-amplification step above, except with 100 µM GAT-COM primers.

### GAM library preparation and high-throughput sequencing

The amplified samples were purified using SPRI beads (0.725 or 1.7 ratio of beads per sample volume), and prepared for sequencing as previously described using the Illumina Nextera XT library preparation kit (Illumina #FC-131-1096) or an in-house library preparation protocol^10^. Following library preparation, the DNA was purified using SPRI beads (1.7 ratio of beads per sample volume) and an equal amount of DNA from each sample was pooled together (up to 196 samples). The final pool was additionally purified three times and analyzed using DNA High Sensitivity on-chip electrophoresis (Agilent 2100 Bioanalyzer). Sequencing was completed using an Illumina NextSeq 500 machine, according to manufacturer’s instructions, using single-end 75 bp reads. The number of sequenced reads for each sample can be found in **Supplementary Table 7**.

### Publicly available GAM datasets

GAM data produced from VTA DNs of animals treated with saline was previously published^10^ and is publicly available in the GEO portal (accession GSE148792) and the 4D Nucleome data portal (https://data.4dnucleome.org)^74^. GAM data produced from pyramidal glutamate neurons (PGNs) from the CA1 region of the hippocampus (HC) and GAM data produced from embryonic stem cells (ESC) was previously published^10,41^ and is publicly available in the GEO portal (accessions GSE64881 and GSE148792) and the 4D Nucleome data portal. Publicly available GAM datasets were downloaded from the 4D Nucleome portal and processed as described below. Sample specifications are listed in the table below.

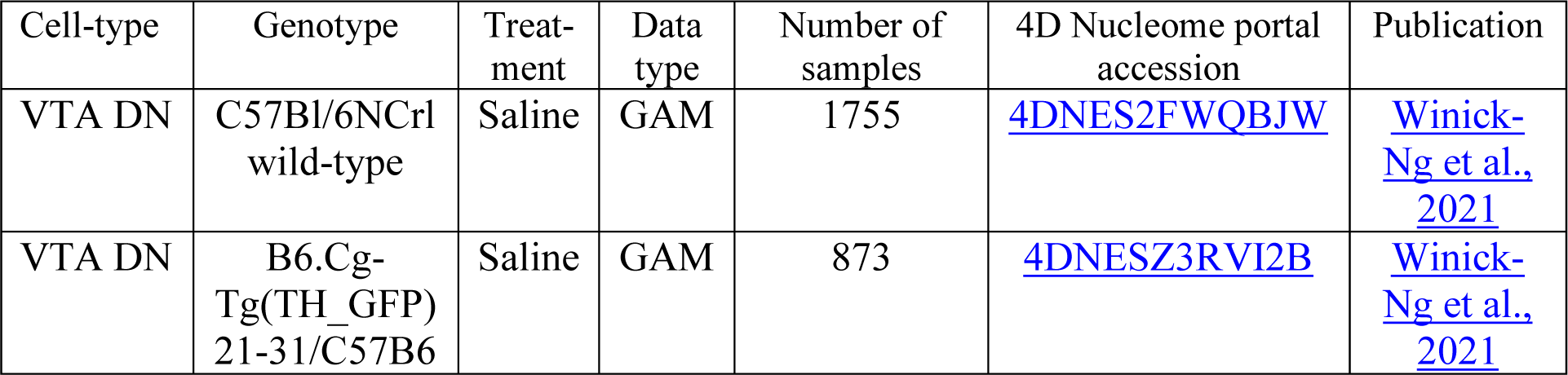
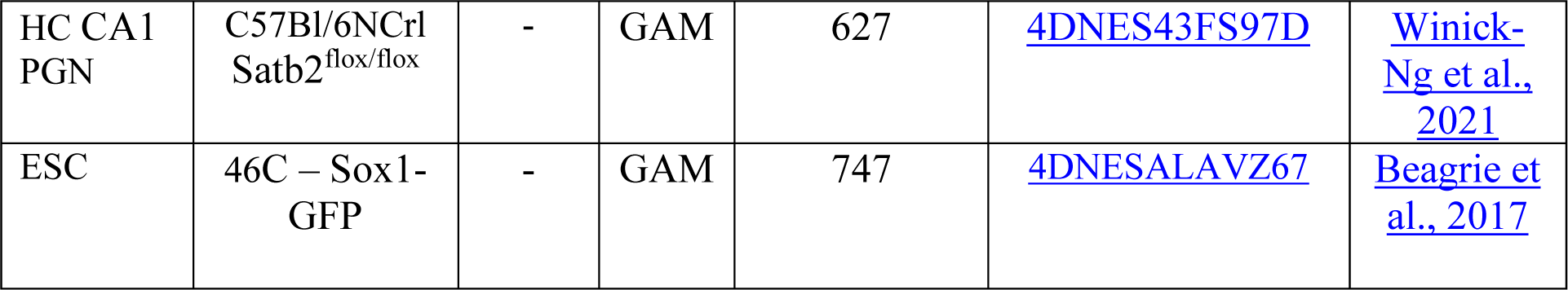

### GAM data sequence alignment

Sequence read alignment was performed as previously described^10,29^. In brief, sequence reads from each GAM library were mapped to the mouse genome assembly GRCm38 (Dec. 2011, mm10) using Bowtie2 with default parameters^75^, and removing reads with mapping quality <20, PCR duplicates and non-uniquely mapped reads.

### GAM data window calling and sample quality control

Positive genomic windows present in each GAM library were identified as previously describe^10,76^. Briefly, the number of nucleotides was calculated for equal-sized genomic windows (250 kb or 40 kb) in each GAM sample. Next, the percentage of orphan windows (i.e. positive windows flanked by both adjacent negative windows) was calculated for every percentile of the nucleotide coverage distribution to identify the percentile with the lowest percent of orphan windows for each GAM sample, and used as the optimal coverage threshold for positive window identification in that sample.

Quality control metrics including the percentage of orphan windows in each sample, number of uniquely mapped reads to the mouse genome, and correlations from cross-well contamination for every sample can be found in **Supplementary Table 7**. Each sample was considered to be of good quality if they had < 70% orphan windows, > 50,000 uniquely mapped reads, and a cross-well contamination score (Jaccard index) of < 0.4. For each treatment and mouse replicate, individual experimental batches were also checked for the same quality control metrics to ensure minimal batch-to-batch variability. In the 1-day cocaine TH-GFP mouse dataset, 34 single NPs were collected, of which 32 passed quality controls. To create *in-silico* 3 NP samples, sequenced reads identified for the 30 single NPs with highest quality scores were combined in random sets of 3 single NPs to create 10 *in-silico* 3NP samples. The 2 remaining NPs were excluded from the final dataset. The number of samples in each treatment passing quality control is summarized in **Extended Data Fig. 1c**.

### Generation of pairwise chromatin contact matrices for visualization

Pairwise contact matrices were generated as previously described^10^, by calculating pointwise mutual information (PMI) for all pairs of windows in the genome, followed by bounding between −1 and 1 to produce a normalized PMI (NPMI) value. For visualization, the genomic regions displayed in each figure are scaled between a range of 0 and the 99th percentile of NPMI values in each treatment.

### Insulation score and topological domain boundary calling

TAD calling was performed by calculating insulation scores in NPMI normalized pairwise GAM contact matrices at 40-kb resolution, as previously described^10,21^. In brief, the average interaction strength of all chromatin contacts within a sliding square of varying size (ranging from 240 - 1040 kb in increments of 80 kb) was calculated for each treatment and replicate, then log normalized relative to all calculated scores of a given square size across each chromosome. Insulation score values of all samples are archived in **Supplementary Table 11** in a permanent data repository^77^.

TAD boundaries were identified using the 400kb-insulation square size. Boundaries overlapping by at least 1 genomic bin (40 kb) were merged, then refined to consider the minimum insulation score within the boundary and one window on each side. 120-kb boundaries separated by at least 1 genomic bin were considered different between datasets (chromosome Y was excluded from this analysis). For comparison of Cocaine 14d-specific boundaries, the union of both replicates were used. TAD border coordinates can be found in **Supplementary Table 4**.

### MELTRONIC analysis

Contact density values were calculated from NPMI normalized pairwise GAM contact matrices at 40-kb resolution with 10 equidistant square sizes between 240 - 1040 kb (in 80 kb increments). Our previously published statistical framework MELTRON^10^ was extended to include the measurement of chromatin condensation, by performance of an additional one-sided Kolmogorov-Smirnov test and the option to perform genome-wide analyses (https://github.com/pombo-lab/MELTRONIC). MELTRONIC compares cumulative probability distributions of contact density values calculated for each input sample and genomic interval and computes the maximum distance between the distributions by applying a Kolmogorov–Smirnov test. Obtained p values were corrected for multiple testing using the Bonferroni method, and –log10 transformed to obtain a melting score.

Melting scores were calculated with MELTRONIC in 120 kb sliding windows across the entire genome by comparing GAM samples of DNs from cocaine treated animals to the saline treated reference data. Genomic regions in which more than 50% of NPMI values were missing within the insulation square, were excluded from the analysis. Melting scores were assigned to the central 40 kb bin and mean smoothed across three genomic bins. Genomic bins with a melting score above 5 or below −5 were identified as ‘melting’ or ‘condensing’, respectively, which identify the 33% of the genome with the most extensive (de)condensation changes, in comparison with 45% when comparing two cell types (**Extended Data Fig. 5c, d**). For genome-wide analysis, regions with reproducible melting states in both cocaine 14d replicates were considered. A list of genome-wide MELTRONIC scores of all comparisons is reported in **Supplementary Table 5**.

### Identification of compartments A and B

Compartments were determined using 250 kb resolution co-segregation matrices, as previously described^10^. Briefly, each chromosome was expressed as a matrix of observed interactions O(i, j) between locus i and locus j, and a matrix of expected interactions, E(i, j), where each genomic window pair represented the average number of contacts with the same distance between i and j. The observed over expected O/E(i, j) matrix was determined by dividing O by E. A correlation matrix, C(i, j), was then generated between column i and column j of the O/E matrix before applying PCA for the first three primary components of matrix C. The component displaying the highest correlation with GC content was extracted, and loci with PCA eigenvector values with the same sign and the strongest correlation with GC content were designated as A compartments, while those with the opposite sign were identified as B compartments. For chromosomes 5, 12 and 14, PC1 was selected based on highest correlation with transcriptional activity, as the PC that correlated most with GC content did not display a typical AB compartmentalization pattern^78^.

Eigenvector values within compartment A on the same chromosome were normalized within the range of 0 to 1, whereas values within compartment B on the same chromosome were normalized within the range of −1 to 0. A full list of eigenvector values and assigned compartment associations can be found in **Supplementary Table 8**.

### MELTRON analysis of long genes

For calculation of melting scores of long genes, MELTRON was applied on protein coding genes > 280 kb in length (n = 574), with the contact density of cocaine samples compared to the saline treated reference. Genes with a melting score above 10 or below −10 in were identified as ‘melting’ or ‘condensing’, respectively. Genes with reproducible melting states in both 14-day cocaine replicates were visualized in Fig. 3a and their transcription states analyzed further. A full list of all melting scores can be found in **Supplementary Table 5**.

### Transcription factor binding site motif analysis at Rbfox1 promoters and a putative cocaine addiction SNP

Transcription factor binding site enrichment (TFBS) analyses were computed for *Rbfox1* promoters and putative SNP +/- 500 bp. Enrichment analyses were performed using two independent methods from the MEME suite programs, ‘Analysis of Motif Enrichment’ (AME)^79^ and ‘Simple Enrichment Analysis’ (SEA)^80^ with default parameters and shuffled input sequences as the control sequences. In both analyses, sequence enrichment was determined using the HOCOMOCO mouse (V11, FULL) database. Sequence coordinates analyzed and a list of enriched transcription factor binding site motifs in both analyses can be found in **Supplementary Table 9**.

### Polymer modeling of the Rbfox1 locus

To investigate the 3D structure of the Rbfox1 locus, we employed the Strings and Binders Switch (SBS) polymer model^81,82^. In the SBS model, a chromatin region is represented as a string of beads incorporating binding sites of different types which can interact with cognate diffusing binding molecules. To infer the SBS polymers for the *Rbfox1* locus in saline- and cocaine-(1 and 14 days) treated VTA DNs, and in untreated pyramidal glutamatergic neurons (PGNs) from the hippocampus^10^, we employed PRISMR, a machine-learning-based method that takes pairwise contact data as input, such as Hi-C^46^ or GAM^83^, and returns the optimal arrangement of binding sites along the polymer to fit the input. Here, we used as input GAM experimental data with NPMI normalization on a 5 Mb region around the *Rbfox1* gene (chr16: 4,800,000 - 9,810,000) at 30-kb resolution in saline and cocaine (1 and 14 days) treated VTA DNs, and in untreated PGNs. As output, PRISMR returned SBS polymers made of 1,670 beads, including 7 different types of binding sites, in all cases.

Next, to generate ensembles of 3D conformations representing the locus folding, we performed standard Molecular Dynamics (MD) simulations of the SBS model for each of the considered cases. In these simulations, the system of beads and binders evolves according to the Langevin equation with classical interaction potentials^84^, with parameters employed in previous studies^10,85^. Specifically, the hard-core repulsion between all beads and binders is modeled with a truncated, shifted Lennard-Jones (LJ) potential. The interactions between beads and cognate binders are modeled by an attractive, short-ranged LJ potential, with an affinity taken in the range from 3.0 to 8.0 K_B_T (where K_B_T is the Boltzmann constant and T the system temperature) and equal for all binding site types for the sake of simplicity. An additional non-specific interaction with lower affinity (from 0 K_B_T to 2.7 K_B_T) is set among the polymer and the binders. Polymers are initially set in self-avoiding conformations and the binders are randomly placed in the simulation box. For the sake of simplicity, beads and binders have the same diameter *s* = 1 and mass m = 1, expressed in dimensionless units. The total binder concentration is taken above the transition threshold to ensure the polymers fold in their equilibrium globular phase^81^. For each of the considered cases, we obtained ensembles of up to 1100 distinct conformations at equilibrium. The MD simulations were performed using the freely available LAMMPS software (v.5june2019)^86^.

To obtain *in-silico* GAM NPMI matrices from the ensembles of 3D conformations, we applied the *in-silico* GAM algorithm^83^. Specifically, we simulated the GAM protocol with 3 NPs per sample by aggregating the content of three *in-silico* slices into one in silico tube^10^, by using 586 tubes for saline, 335 tubes for cocaine 1 day, and 404 tubes for cocaine 14 day treatments, as well as 209 tubes for untreated PGNs, as in the corresponding GAM experiments. Finally, we applied the NPMI normalization. To compare *in-silico* against experimental NPMI GAM matrices, we computed Pearson’s correlation coefficients.

To quantify the changes in 3D chromatin organization in the different considered cases, we measured spatial distances between different locations of interest within the *Rbfox1* gene in the model 3D structures. To convert distances from dimensionless units σ to physical units (nm), we estimated the physical diameter of the bead σ by optimizing the similarity between the *in-silico*^83^ and experimental NPMI GAM matrices in the different cases. We obtained values for σ of 43 nm in saline, 49 nm in cocaine 1 day, 56 nm in cocaine 14 days and in untreated PGNs, that corresponds to a chromatin compaction factor comprised between 54 bp/nm and 70 bp/nm, consistent with values found in literature^87–89^ and used in previous polymer modeling studies^43,46,83^. To visualize chromatin organization in the different cases, we also rendered example 3D structures from the derived ensembles by performing a third-order spline of the polymer bead positions. We highlighted the *Rbfox1* gene region in blue color, its left/right flanking regions in dark/light gray respectively and represented different locations within the gene with colored spheres. Analyses and plots were produced with the Anaconda package v.22.9.0 and the rendering of 3D structures was produced using POV Ray, v.3.7 (http://www.povray.org/). All polymer model 3D structures and pairwise distances produced for the analyses of this work are archived in **Supplementary Tables 12-16** in a permanent data repository^77^.

### Determining differential contacts and hotspots between GAM datasets

Significant differences in pairwise contacts between two GAM datasets was determined as previously described^10^. Briefly, genomic windows with low detection (< 2% of the distribution of all detected windows in each chromosome) were removed from both datasets, and NPMI contact frequencies were normalized by computing the Z-score transformation. A differential matrix D was computed by subtracting the two normalized matrices and a 5-Mb distance threshold was applied. The top 5% significant differential contacts were obtained by fitting a normal distribution curve for each chromosome and determining the upper and lower 5% from the curve.

Next, preferred (hotspot) contact regions for each compared dataset were determined, as previously described^29^, by first quantifying the number of top 5% significant contacts in each genomic window for each dataset. A ‘hotspot score’ was determined by computing the difference between the number of significant contacts in each genomic window for each dataset. Significant hotspot scores were obtained by fitting a normal distribution curve for each chromosome and determining the upper and lower 5% from the fitted curve. A full list of genome-wide hotspot scores can be found in **Supplementary Table 6**.

### Gene ontology and synaptic gene ontology enrichment analysis

Gene ontology (GO) term enrichment analysis was performed using WebGestalt over-representation analysis (ORA)^90^ using the ‘Geneontology’ functional database category and ‘Biological Process’ as function database name. Overlap of expressed genes with 3D genome features (TAD boundaries, melting/condensing regions, compartments, hotspots) were determined with the valr R package v.0.6.8^91^, bedtools v2.30.0^76^ or pybedtools^92^. All DN expressed genes were used as the background dataset. Default parameters were used to determine enrichments and the top 20 terms were reported. For **Fig. 2b** and **Extended Data Fig. 4a**, example GO terms with non-redundant gene identifiers and a significant enrichment (p-adj < 0.05) were selected. A full list of unfiltered GO term enrichment results can be found in **Supplementary Table 10.**

For synaptic gene ontology (SynGO) analysis^33^, mouse gene ids were converted to human gene homologs with the biomaRt R package v.2.55.2^93^. DN expressed genes were used as the background dataset and enrichments computed with the default parameters of the SynGO release 1.1 (20210225 release). Biological process terms that were significantly enriched (p-adj < 0.05) in at least one set were visualized in **Fig. 2c, Extended Data Fig. 4b** and **Extended Data Fig. 5f**. A full list of unfiltered SynGO term enrichment results can be found in **Supplementary Table 10.**

### Collection of single nucleotide variants underlying substance addiction

Single nucleotide polymorphisms (SNPs) associated with cocaine dependence and comorbid addictions to alcohol, nicotine and opioids were collected from publicly available resources^19,24,94^. In case of positional overlap, SNP with lowest p-value was used before conversion of genome coordinates of noncoding SNPs from human to mouse coordinates with the liftOver R package v.1.22.0. Conservation in the genomic neighborhood of the SNPs was assessed with CNEr v.1.34.0^95^. SNPs found in genomic regions in which at least 5 out of 7 bps are conserved were considered for downstream analyses. A full list of addiction SNPs can be found in **Supplementary Table 1**.

### Collection of immediate early genes and genes associated with cocaine addiction

The gene set for immediate-early gene (IEG) induction in rodent brain was obtained by intersecting datasets of previously identified neural gene expression signatures in response to acute cocaine^96,97^ or kainic acid^98^. The cutoff for differentially expressed genes was adjusted for each dataset. In bulk tissue RNA-seq of rat brain, genes with adjusted *P* value < 0.05 in the comparison between “Cocaine Challenge” (SC) (10 mg/kg i.p; 1h after cocaine) and “No Challenge” (S24) were considered as differentially expressed^96^. To identify the transcriptional response in the mouse nucleus accumbens to acute cocaine (20 mg/kg i.p; 1h after cocaine) in single nucleus RNA-seq data, a combined cutoff for significance (p-value attached < 0.05) and log2 fold change (log2 fold change > 0.5) was used ^97^. In RNA-seq data of mouse hippocampal neuronal nuclei (nuRNA-seq), genes with adjusted p value less than 0.05 and log2 fold-change of at least 1 were considered as differentially expressed after systemic administration of kainic acid (25 mg/kg i.p; 1h after kainic acid)^98^. Genes that were significantly upregulated in at least two datasets comprised the final IEGs gene set (141 genes). The genomic coordinates of immediate early genes are provided in **Supplementary Table 1**. Expression levels of immediate early genes and fold changes after cocaine exposure can be found in **Supplementary Table 3**. Genes associated with cocaine addiction were collected from publicly available resources^19,20,57,58,62,99,100^. The genomic coordinates of cocaine addiction associated genes and are provided in **Supplementary Table 1**. Expression levels of cocaine addiction associated genes in dopaminergic neurons and fold changes after cocaine exposure can be found in **Supplementary Table 3**.

### Isolation of the VTA for single-nucleus RNA-sequencing

Mice were anaesthetised with 4% isoflurane and swiftly decapitated. Brains were removed and briefly washed in ice-cold sterile PBS, they were placed on fresh filter paper and a block of tissue containing the VTA was dissected using a sterile razor blade. The tissue was then snap-frozen in isopentane (2-Methylbutane; Sigma-Aldrich) at −55°C.

### Isolation of nuclei for single-nucleus RNA-sequencing

VTA nuclei were isolated and prepared for single-nucleus RNA-sequencing as previously described^17^. VTA tissue was removed from the −80 °C and transferred into a 1 ml Dounce homogenizer containing 1 ml of pre-chilled homogenization buffer (250 mM sucrose, 25 mM KCl, 5 mM MgCl_2_, 10 mM Tris pH 8.0, 1 mM DTT, 1x protease inhibitor (Roche, 11873580001), 0.4 U/ul RNAse inhibitor (Takara, 2313B), 0.2 U/µl SUPERase•In (Invitrogen, AM2696), 0.1% Triton X-100). Tissue was treated with 5 strokes of the loose pestle, followed by 15 strokes of the tight pestle and then filtered through a 40 µm cell strainer. Nuclei were spun down at 1000 x *g* for 8 min at 4 °C. The pellet was then resuspended in a homogenization buffer with the final volume of 250 µl on ice. The suspension was mixed with 250 µl of 50% iodixanol solution (25 mM KCl, 5 mM MgCl_2_, 10 mM Tris (pH 8.0), 50% iodixanol (60% stock from STEMCELL Technologies, 7820), 1× protease inhibitor (Roche, 11873580001), RNase inhibitor (0.4 U/µl; Takara, 2313B), SUPERase·In (0.2 U/µl; Invitrogen, AM2696), and 1 mM DTT) and overlaid on top of 29% iodixanol solution (25 mM KCl, 5 mM MgCl_2_, 10 mM Tris (pH 8.0), 29% iodixanol, 1× protease inhibitor (Roche, 11873580001), RNase inhibitor (0.4 U/µl; Takara, 2313B), SUPERase·In (0.2 U/µl; Invitrogen, AM2696), and 1 mM DTT) in an ultracentrifuge tube (Beckman Coulter, 343778) on ice. Gradients were spun down in the ultracentrifuge (Beckman Coulter, MAX-XP) using a swing bucket rotor (Beckman Coulter, TLS 55) at 14,000 *g*_max_ (∼10900 *g*_average_) for 22 min at 4°C with slow acceleration and deceleration. Supernatant was carefully removed, and pellets were resuspended in ice-cold bovine serum albumin (BSA) blocking solution (1× phosphate-buffered saline (PBS) (AM9625, Ambion), 0.5% BSA (VWR, 0332-25G), 1 mM DTT, 2.4 mM MgCl_2_, and RNase inhibitor (0.2 U/µl; Takara, 2313B)) and incubated on ice for 15 min. Before staining, splits were taken for the controls (isotype control, negative control, 7AAD only control, NeuN only control). The neuronal marker NeuN antibody (1 µg/µl, 1:5670, Millipore, MAB3777X) was added to the sample and NeuN-only control. The control antibody (0.2 µg/µl, 1:1134, STEMCELL Technologies, 60070AD) was added to the isotype control. Antibodies were incubated for 10 min at 4 °C in the dark. After incubation, 1 ml of BSA blocking buffer was added and centrifuged at 1000 x *g* for 10 min at 4 °C in a swing bucket. Pellets were resuspended in 200 µl BSA blocking buffer and filtered through a 35 µm strainer. Samples were then filled with BSA blocking buffer to a total volume of 500 µl and 0.75 µl of 7AAD (1 mg/ml, Sigma) was added.

FACS was performed using BD FACSAria III sorter using a 75 µm nozzle and controlled by BD FACSDiva 8.0.1 software. Single color controls were used for compensation. Gates were set based on the FACS controls. Nuclei were selected using a FSC-A/SSC-A gate, doublets were removed using FSC-W/FSC-H and SSC-W/ SSC-H gates, nuclei were then further selected on the basis of 7AAD staining, and neuronal nuclei were sorted on the basis of NeuN staining (**Extended Data Fig. 1e**). Nuclei were collected into 5 µl of BSA blocking buffer at 4 °C and directly processed for 10x Genomics library preparation.

### Single-nucleus mRNA library preparation

The chromium Single Cell 3’ Reagent kit v3 (10x Genomics, 1000075) was utilized for most library preparations, with the standard protocol applied. For the library preparation of four biological replicates, the chromium Single Cell 3’ Reagent kit v2 (10x Genomics, 1000009) was used (see **Extended Data Fig. 1f**). In brief, nuclei were counted under a brightfield microscope and mixed with the reverse transcription mix. Gel Beads were added and the mix was partitioned on Chips B (10x Genomics, 1000073) into GEMs using the Chromium Controller (10x Genomics, PN-120223) for reverse transcription. After reverse transcription, samples were frozen at −20 °C until further processing. Next, cDNA was cleaned, pre-amplified (12 PCR cycles), cleaned with SPRIselect beads and quantified before being frozen again at −20°C. The same quantity of cDNA was used during fragmentation, end-repair, and A-tailing for most samples. Fragments were then cleaned up using SPRIselect reagent and processed through the steps of adapter ligation, SPRIselect cleanup, and sample index PCR (using Chromium i7 Sample Indices (10x Genomics, PN-120262) for 11 PCR cycles). Libraries were cleaned up with SPRIselect reagent and quantified using the Qubit HS dsDNA Assay Kit (Thermo Fisher Scientific, Q32854) with a Qubit Fluorometer, and also using the High Sensitivity DNA Kit (Agilent, 5067-4626) with an Agilent 2100 Bioanalyzer. Libraries were pooled according to the expected amount of nuclei per sample and sequenced using an Illumina HiSeq 2000 machine according to manufacturer’s instructions.

### Single-nucleus RNA-seq data processing: mapping, expression, and QC

Raw RNA sequencing data was processed using the pigx-scrnaseq pipeline, version 1.1.7^101^. In short, the sequencing reads were mapped using STAR^102^ on the mm10 version of the mouse genome. The digital expression matrix was constructed using the mm10 mouse gene annotation GRCm38.82, downloaded from the ENSEMBL database^103^. Gene expression was quantified by counting the reads overlapping complete gene models (both exons and introns) and normalized using the Seurat pipeline with default parameters. Ambient RNA was removed from the digital expression matrix using CellBender^18^ with the default parameters.

### Single-nucleus RNA-seq quality control

Putative droplet doublets were detected using scDblFinder^104^ and only singlet cells were kept for further analysis. To ensure high cell quality, cells with fewer than 2000 detected features were removed from the analysis. In addition, cells with a ratio of exonic to intronic reads lower than the 5% percentile or greater than the 95% percentile (calculated on a per sample basis) were also removed from further analysis. Because the previous stringent quality control filters removed the majority of cells sequenced using 10x v2 chemistry (saline, and cocaine day 1 replicates 1 and 2), the v2 chemistry samples were removed from any further analysis.

The resulting filtered data was integrated using Conos^105^. Integrated data processed using Seurat^106^ - data was normalized, scaled, transformed using principal component analysis. Data was embedded in a low dimensional state using the UMAP algorithm^107^.

Clustering was completed using the FindClusters method from the Seurat package using the Conos derived cell distance graph. The clustering was obtained using the Louvain algorithm with resolution parameter set to 0.1.

### Single nucleus RNA-seq dopaminergic cell identification

Cells were scored with a set of well-known dopaminergic marker genes *Th, Slc6a3, Nr4a2, Slc18a2, Snca, Foxa2, Lmx1b, Kcnj6*^26^, using the Seurat function AddModuleScore. Cells belonging to the cluster with the highest median dopaminergic score were regarded as putative dopaminergic neurons. To remove nuclei which may have come from the substantia nigra (SN), the DN-containing region neighbouring the VTA, a set of known SN markers *Sox6*, *Aldh1a7*, *Ndnf*, *Serpine2*, *Rbp4*, *Fgf20* were used as an input to the AddModuleScore function, and all cells with a score greater than 0.65 were removed from the analysis. The resulting DNs were processed using the default Seurat pipeline, including read count normalization, scaling, and PCA calculation. The data was then embedded in 2D space using the UMAP algorithm. Cluster specific markers were determined using the FindAllMarkers function from the Seurat package.

To obtain a robust set of markers specific for the “IEG” cluster (**Fig. 5c**), differential genes were iteratively defined by comparing the IEG cluster with all other clusters, and taking genes which were significantly enriched in the IEG cluster in all comparisons.

Gene set scores for melting genes, hotspot genes and TAD boundary genes were obtained using the AUCell function from the AUCell bioconductor package^108^.

### Single-nucleus RNA-seq differential expression analysis

Differential expression analysis was calculated for the complete datasets of dopaminergic cells, and separately for the IEG subcluster. The per cell count data was summarized as pseudo-bulk values using the muscat package^109^. Differential expression analysis was then completed using all three of the available methods implemented in the muscat package: edgeR, DESeq2 and limma. None of the methods were sensitive enough to detect individual differentially expressed genes with an FDR-adjusted P value lower than 0.05. A full list of expression values and differential expression analysis results can be found in **Supplementary Table 3**.

Gene set differential expression was performed by first computing the mean and median fold change of IEGs or addiction genes in both the complete DN dataset and the IEG cluster. To determine the significance of the mean (and median) fold change, the mean (median) fold change was calculated for 1000 random subsets of genes of the same size as the corresponding gene sets (**Extended Data Fig. 7g**). The Z-score and *P* value was calculated by comparing the measured fold change of the true corresponding gene set with the distribution of the random permutations.

### Integration of spatial transcriptomics MERFISH data

The single-cell MERFISH (scMERFISH) spatial transcriptomics dataset was downloaded from the Allen brain atlas server^27^, along with the corresponding cell and cluster annotations specified below.

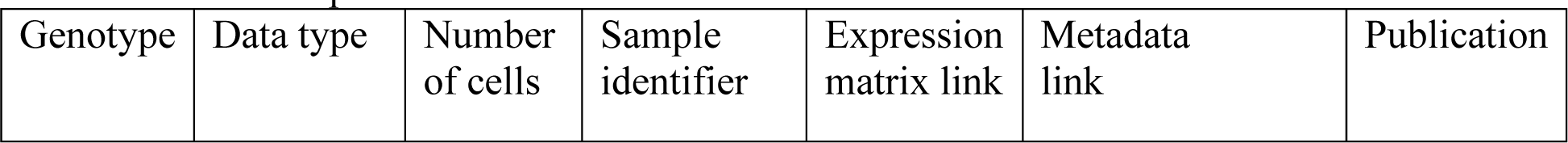
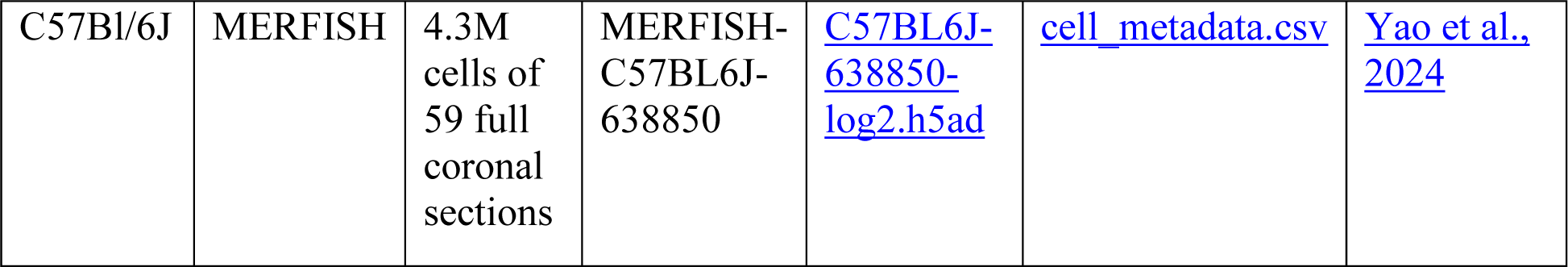

Cells belonging to the VTA were selected from the dataset by taking a subset of cells with the “MB Dopa” class annotation. Cells belonging to the substantia nigra part of the midbrain were filtered based on the high expression of the following markers: *Ndnf*, *Tlll1*, *Epha4*, *Rgs8*. To compare the scMERFISH and snRNA-seq data, both datasets were summarized to the cluster level by taking the average expression of all genes in all corresponding clusters. Next, genes which had linear covariation in both datasets were selected by calculating the variance separately for the scMERFISH and snRNA-seq data and choosing genes with a log2 ratio of (snRNA-seq variance) / (scMERFISH variance) between −0.25 and 0.25 for further analysis. Pearson correlation coefficients were calculated between each snRNA-seq and scMERFISH cluster, and the resulting matrix was visualized using the ComplexHeatmap^110^ package. The correspondence between snRNAseq and scMERFISH clusters was determined by selecting the pair of clusters with the largest calculated correlation coefficient.

## Data availability

Raw fastq sequencing files for GAM datasets generated for this manuscript, together with non-normalized co-segregation matrices, normalized pair-wised chromatin contacts maps and raw GAM segregation tables are available from the GEO repository under accession number GSE254508 and from the 4DN data portal (https://data.4dnucleome.org/) under accession identifiers 4DNESYLI75YL (1 day cocaine, wild-type), 4DNESMAQEPWU (1 day cocaine, TH-GFP), and 4DNESVF6WL86 (14 days cocaine, wild-type). Raw fastq sequencing files for GAM datasets of VTA DNs from animals treated with saline are available from the 4DN data portal under accession identifiers 4DNES2FWQBJW (wild-type) and [4DNESZ3RVI2B](https://data.4dnucleome.org/experiment-set-replicates/4DNES2FWQBJW/) (TH-GFP). Insulation score values of all samples are archived in table S11 in a permanent data repository^77^. All polymer model 3D structures and pairwise distances produced for the analyses of this work are archived in table S12-16 in a permanent data repository^77^.

Raw fastq single nucleus RNA sequencing files, bigwig tracks of DNs and count matrices are available from the GEO repository under accession number GSE254509. Seurat objects of all sequenced neurons (NeuN^+^) and all profiled dopaminergic neurons are archived in a permanent data repository^77^. An interactive application for exploration of snRNA-seq data of dopaminergic neurons is available under the following link: https://shiny.mdc-berlin.de/APombo_VTA. scMERFISH spatial transcriptomics data of dopaminergic neurons of the ventral tegmental area are archived in table S17 in a permanent data repository^77^. Interactive 3D plots indicating locations of IEG expressing DNs and Vip^+^ DNs in the VTA are archived in interactive plot S1-S2 in a permanent data repository^77^. A public UCSC genome browser session with all data produced is accessible under the following link: http://genome-euro.ucsc.edu/s/Kjmorris/GAMcocaine_2024_publicSession.

## Code availability

Processing, analysis and plotting scripts for insulation score calculation and MELTRONIC analyses are available at: https://github.com/pombo-lab/MELTRONIC/.

## Extended Data Figures

**Extended Data Figure 1.**
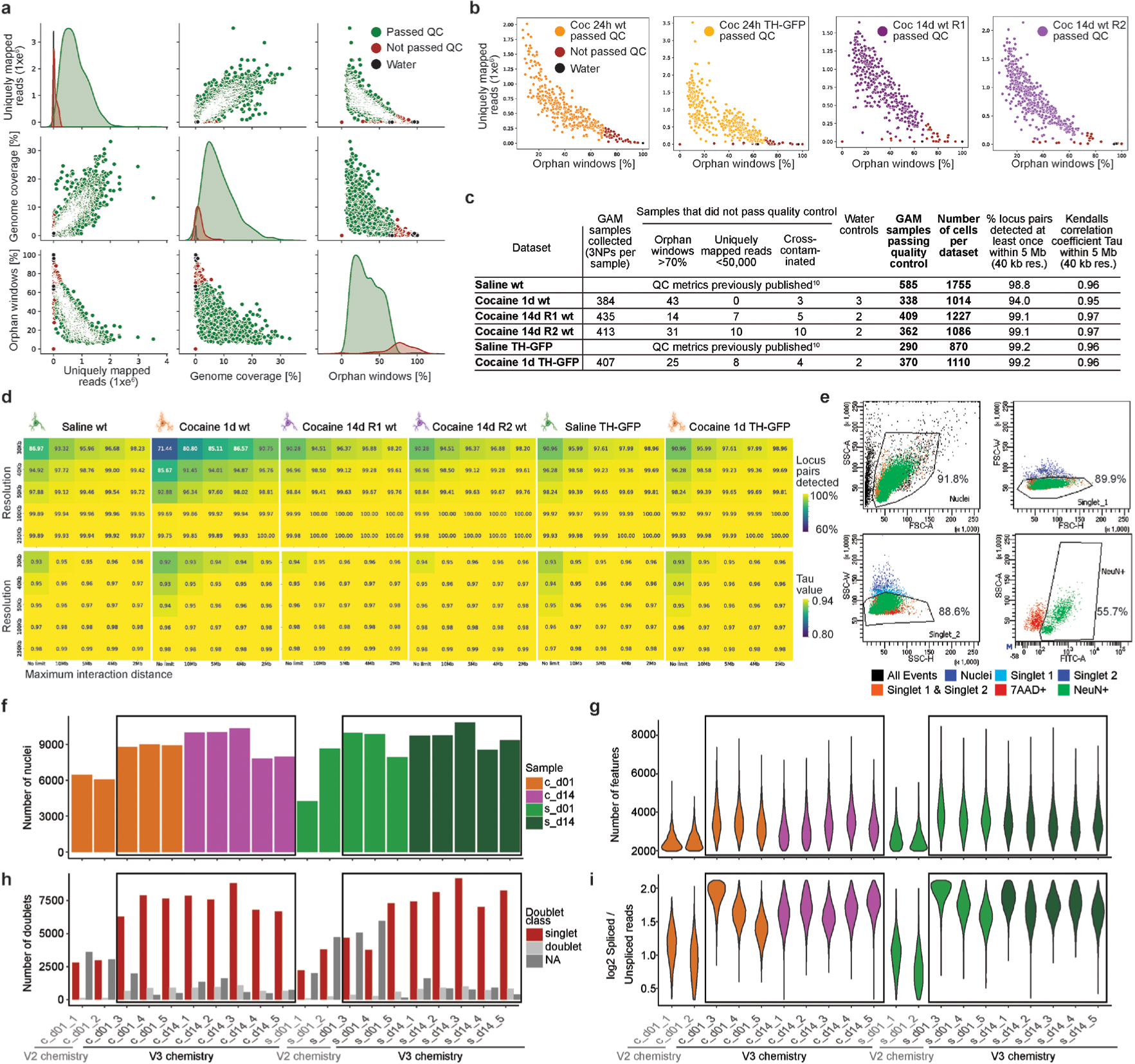
Quality control of immunoGAM and single-nucleus RNA-seq data. **a,** Quality control (QC) measurements (uniquely mapped reads, genome coverage, percentage of orphan windows) for all combined GAM samples collected from ventral tegmental area (VTA) dopamine neurons (DNs). Each data point represents a GAM sample: Green, sample passed QC; Red, sample did not pass QC; Black, water control. **b,** Similar to **a** but shown separately for each dataset produced in this study (number of uniquely mapped reads and percentage of orphan windows). **c**, Summary of GAM data used in this study. All data was collected from 8-week-old mice, 1 or 14 days following an intraperitoneal injection of cocaine (15 mg/kg). GAM data from saline-treated animals were littermates with cocaine-treated animals and was previously published^10^. **d**, Percentage of loci-pairs detected at least once, as well as Kendall’s τ (Tau) coefficients for different GAM resolutions (range 30 - 250 kb) and a range of distance cutoffs (2 Mb - chromosome-wide). We considered a Tau value > 0.95 as an appropriate resolution for a given genomic distance. **e,** Flow cytometry data of a representative sample showing selection criteria for gating of neuronal cells using NeuN (RBFOX-3) as a pan-neuronal marker. **f,** Total number of sequenced nuclei per library. **g,** Distribution of number of detected features (genes) per library. **h,** Number of nuclei classified as singlet / doublet / undefined by the scDblFinder^104^ doublet detection algorithm. **i,** Distribution of the log 2 ratio of exonic to intronic reads for each detected feature in each sample. In panels **f** - **i** cDNA libraries from biological replicates 1 and 2 from 1 day saline and cocaine treatments were prepared using the 10x chromium single cell V2 chemistry, while all other samples were prepared with V3 chemistry (V3 samples are shown inside the black-bounded box). Our conclusion was that the replicates produced with V2 chemistry were of lower quality, and therefore removed from downstream analyses.

**Extended Data Figure 2.**
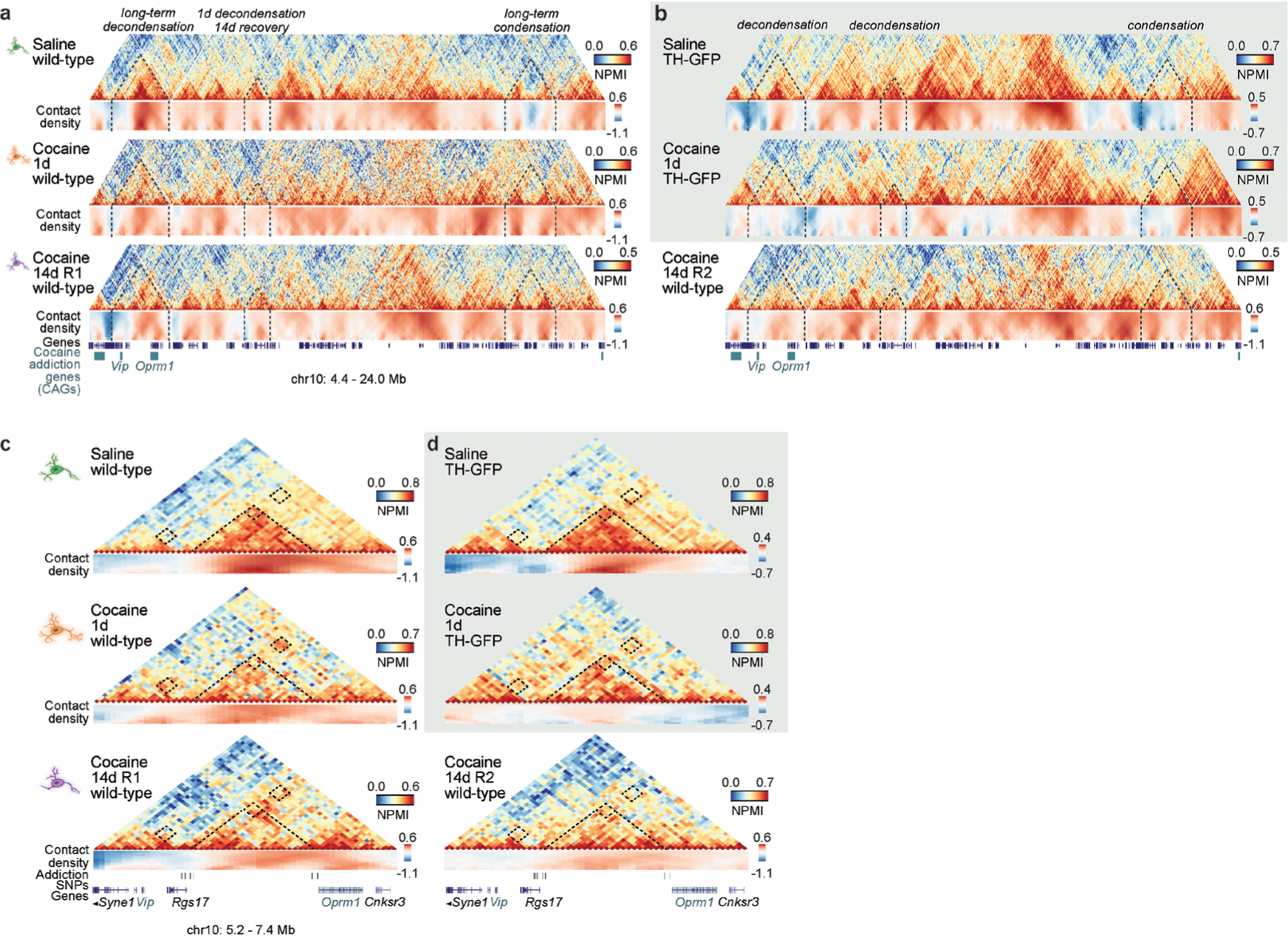
Cocaine induced large-scale disruption of 3D genome structure in GAM replicates. **a,** Similar to **Fig. 1c** but with both cocaine 14 day GAM replicates shown side-by-side. Example of long-term cocaine-induced 3D genome reorganization (chr10: 4.4 - 24 Mb; 40 kb windows). NPMI, normalized pointwise mutual information. Contact density heatmaps show insulation scores ranging from 240-1040 kb. **b,** GAM matrices produced from VTA DNs in TH-GFP animals display similar condensation and decondensation of the highlighted loci 1d after cocaine exposure, though broad contact loss is not as pronounced as in wild-type animals. **c,** Similar to **Fig. 1d** but with both cocaine 14d replicates shown side-by-side. Example of cocaine-induced disruption and rewiring of local contacts seen in both 14-day replicates (chr10: 5.2 - 4.4 Mb; 40 kb resolution). **d,** Similar interaction patterns and rewiring of chromatin contacts after cocaine exposure are observed in GAM data produced from VTA DNs of TH-GFP animals.

**Extended Data Figure 3.**
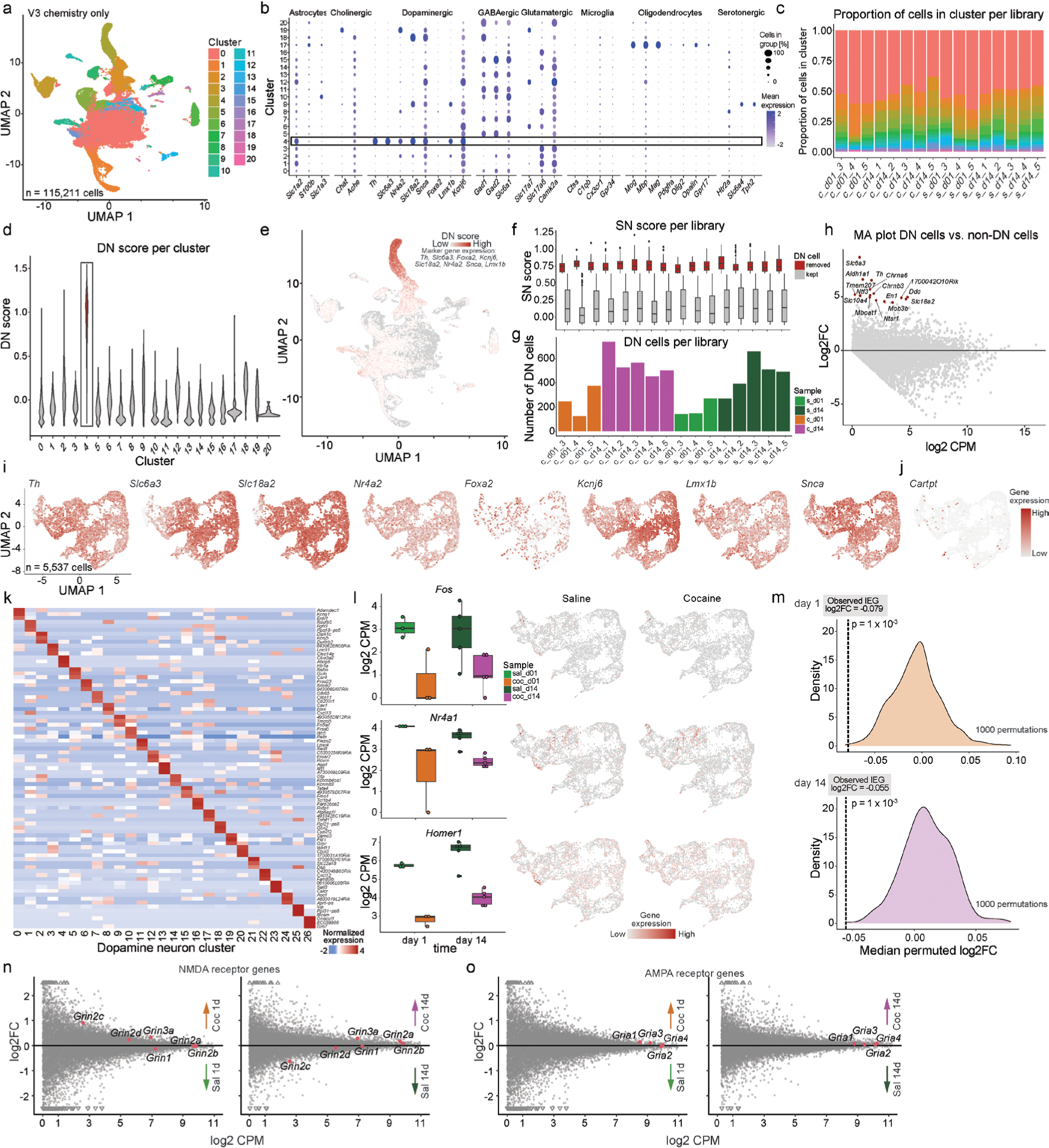
Identification and characterization of VTA dopamine neurons. **a,** UMAP representation of the complete dataset integrated using the Conos algorithm^105^. Sample libraries prepared with 10x chromium single cell V2 chemistry (see **Extended Data Fig. 1f-i**) were excluded from the integration and downstream analyses. **b,** Expression of brain cell type marker genes in the integrated clusters. c, Relative abundance of cell-types captured in each library. **d,** Distribution of the DN gene set score, in each cluster, of a set of known dopaminergic specific genes obtained using the AddModuleScore function from the Seurat package. Nuclei belonging to cluster 4 were classified as putative dopaminergic nuclei (grey box). **e,** Dopaminergic gene set score overlaid on top of the UMAP representation. **f,** Distribution of gene set score of known substantia nigra (SN) markers in the putative dopaminergic nuclei in each sample. Cells with a SN score greater than 0.65 were classified as putative SN cells and removed from further analyses. **g,** Number of dopaminergic nuclei per sample. **h,** MA plot showing the average expression and log2 fold change of VTA DNs versus non-DN cells. Top genes differentially upregulated in dopaminergic cells are well known DN markers *Th*, *Aldh1a1*, and *En1*^111–113^. **i,** Normalized gene expression of known dopaminergic marker genes overlaid on the UMAP representation. **j,** Normalized expression of cocaine response gene *Cartpt*. **k,** Heatmap showing the scaled average gene expression of the top three marker genes in each of the VTA dopaminergic clusters. **l,** Example expression of a selection of IEGs (*Fos*, *Nr4a1*, *Homer1*), related to **Fig. 1g**. Each point shows the average log2 DN pseudobulk transcription of the gene in one replicate of the corresponding condition. The related integrated DN UMAPs represent the expression of the IEG genes in the saline and cocaine conditions for each gene. **m,** Results of the gene set permutation analysis. The density plot shows the distribution of the median fold changes of equal sized randomized gene sets. The dotted vertical line shows the median fold change of the IEG gene set (n = 139). **n and o,** MA plots showing the log2 fold change ratio of cocaine and saline pseudo-bulk RNAseq samples versus the average expression of each gene. Colored dots indicate the average transcription level and the fold change of NMDA receptors in **n** and AMPA receptors in **0**. For **f** and **l**, boxplot whisker length represents 1.5 times the interquartile range.

**Extended Data Figure 4.**
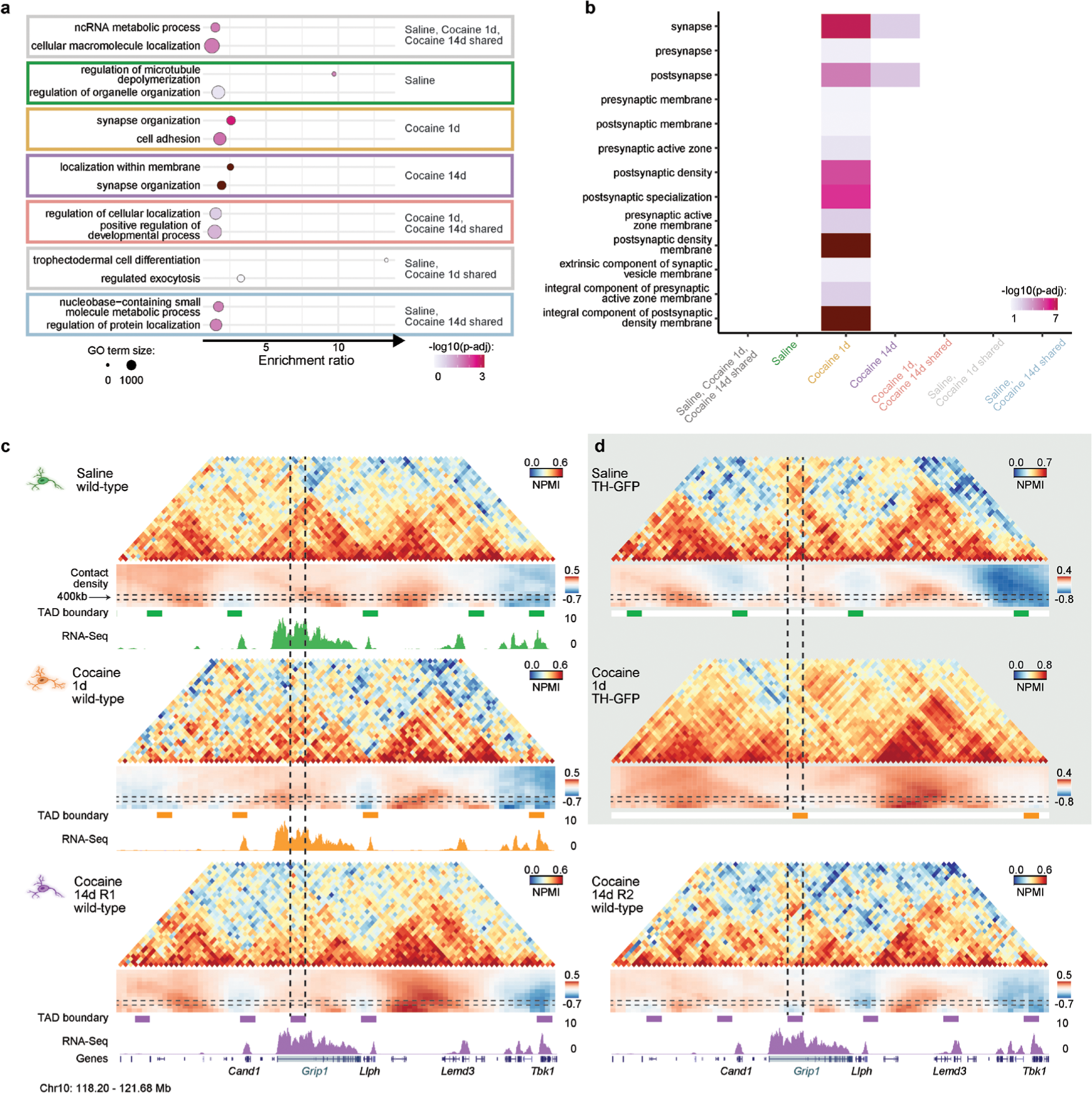
Cocaine-induced TAD boundaries are enriched for synapse related genes. **a,** Similar to **Fig. 2b** but for all groups found in **Fig. 2a**. A complete list of gene ontology (GO) terms can be found in **Supplementary Table 10**. **b,** Similar to **Fig. 2c** but for all groups found in **Fig. 2a**. Synaptic gene ontology (SynGO) enrichments were only found for boundaries specific to 1 or 14 days after cocaine. A complete list of synGO terms can be found in **Supplementary Table 10**. **c,** Similar to **Fig. 2d** but with both cocaine 14d GAM replicates shown side-by-side. *Grip1* overlaps a 14-day specific boundary in both biological replicates (coloured boxes below contact density heatmap; chr10: 118 – 122 Mb). **d,** *Grip1* locus in GAM data produced from VTA DNs in TH-GFP animals. The saline-treated mouse with TH-GFP background is similar to wild-type. However, contact reorganization 1 day after cocaine result in a new TAD boundary at the *Grip1* locus in the TH-GFP animal, not seen in the wildtype mouse.

**Extended Data Figure 5.**
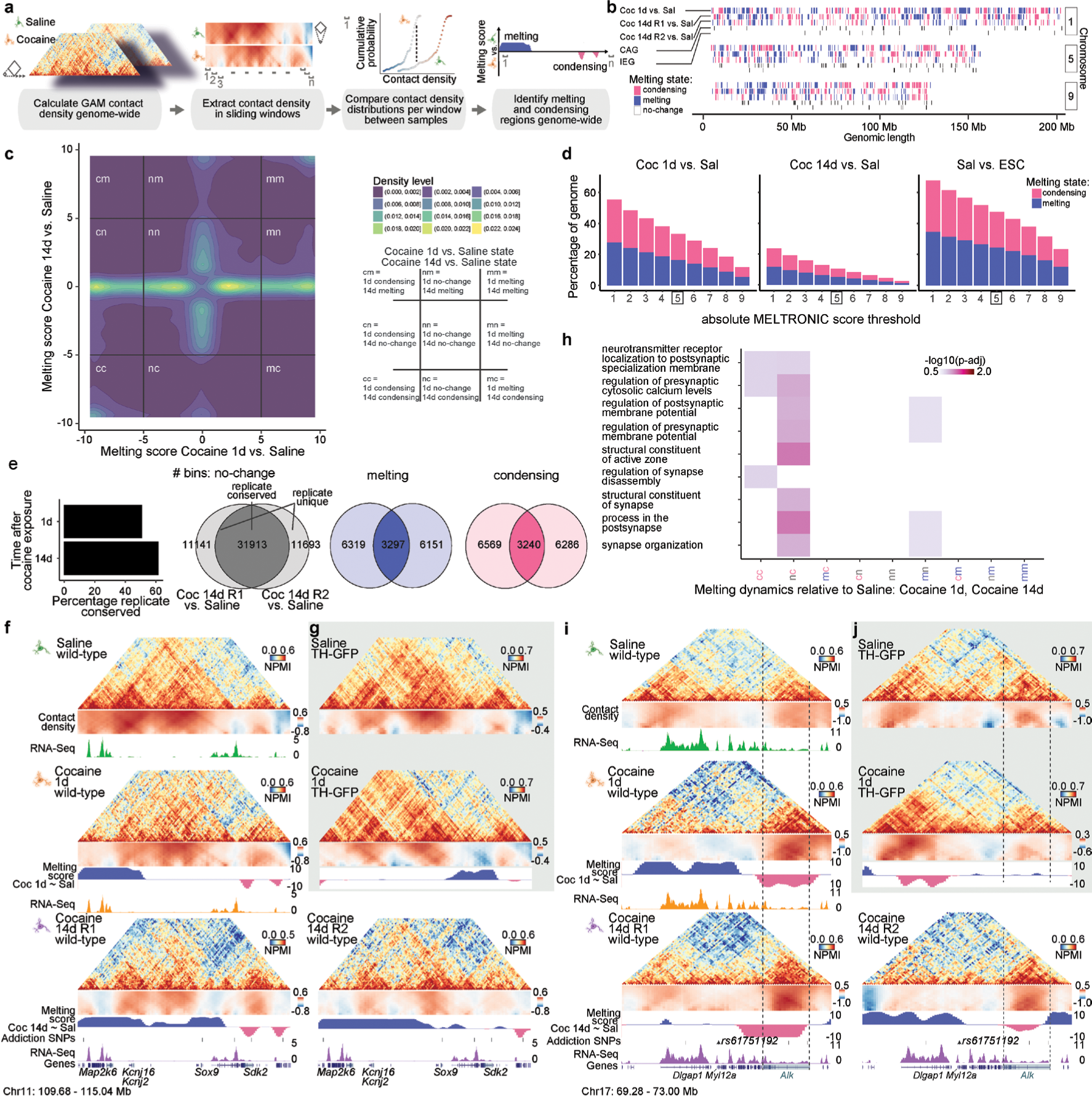
MELTRONIC discovers extensive chromatin melting and condensing after a single cocaine exposure genome-wide. **a,** The MELTRONIC pipeline was applied genome-wide between cocaine- and saline-treated DNs. Contact density value distributions (ranging 240 - 1040 kb) of cocaine-treated DNs were compared to saline-treated DNs in 120 kb sliding windows with a one-sided Kolmogorov–Smirnov test, and visualized as a cumulative probability distribution function. 40 kb genomic bins with a melting score > 5 or < −5 were identified as melting or condensing, respectively (corresponding to a multiple testing corrected *P* < 1 × 10^−5^). **b,** Linear version of **Fig. 2e** of representative example chromosomes. Condensing (pink, melting score < −5), melting (blue, melting score > 5) and non-changing (white) regions in cocaine 1 day vs. Saline (top row), cocaine 14 days R1 vs. saline (2nd row), and cocaine 14 days R2 vs. Saline (3rd row) comparisons. The positions of cocaine addiction genes (CAGs) and immediate early genes (IEGs) are shown below. **c,** Two-dimensional representation of melting score densities in replicate-reproducible bins from the cocaine 14 day R1 vs. saline comparison (y-axis) and cocaine 1 day vs. saline comparison (x-axis). Melting/condensing thresholds (5 and −5, respectively) are indicated as solid black lines. For visualization purposes, bins with melting scores > −1 and <1 in both comparisons were removed. **d,** Assessment of melting score thresholds in the cocaine 1 day vs. saline comparison (left), cocaine 14 day vs. saline (middle; for bins where melting or condensing was reproducible in both replicates), and saline-treated DNs vs. ESCs. The percentage of the genome identified as melting or condensing decreases near linearly with the threshold. The percentage of melting/condensing regions identified by a given threshold is higher between cell types than between treatments. **e**, Histogram (left) showing the percentage of genomic regions found with the same melting state in the cocaine replicates. Venn plots (right) show the replicate overlap for non-melting, melting, and condensing regions. **f,** Similar to **Fig. 2f** but with both cocaine 14-day GAM replicates shown side-by-side, showing melting of *Kcnj16* and *Kcnj2*, and condensing downstream of *Sox9*, in both 14 day replicates (chr11: 110 - 115 Mb). **g,** The *Kcnj16/Kcnj16* locus is similar in a saline-treated mouse with TH-GFP background. In contrast with the wildtype mouse, melting of the *Kcnj2*/*Kcnj16* genes is not detected in the TH-GFP background mice, while the region downstream of *Sox9* has both melting and condensing regions. **h,** SynGO enrichments (p-adj < 0.05) of biological processes were identified for genes encoded in short-term melting and long-term condensing regions. A complete list of synGO terms can be found in **Supplementary Table 10**. **i,** Long-term condensation of the *Alk* gene both 1 and 14 days after a single cocaine exposure (chr17:69.28-73.00 Mb). *Alk* condenses in both 14 day replicates, though more distinctly in R1. The upstream region, containing a putative cocaine addiction SNP (rs61751192) and the *Dlgap1* gene, melts 1 day after exposure, then returns to its pre-melted state in cocaine 14-day R1, but not R2. **j,** Similar to wildtype, the *Alk* region condenses, while the cocaine addiction SNP and the *Dlgap1* region melts, 1 day following cocaine in the TH-GFP mouse.

**Extended Data Figure 6.**
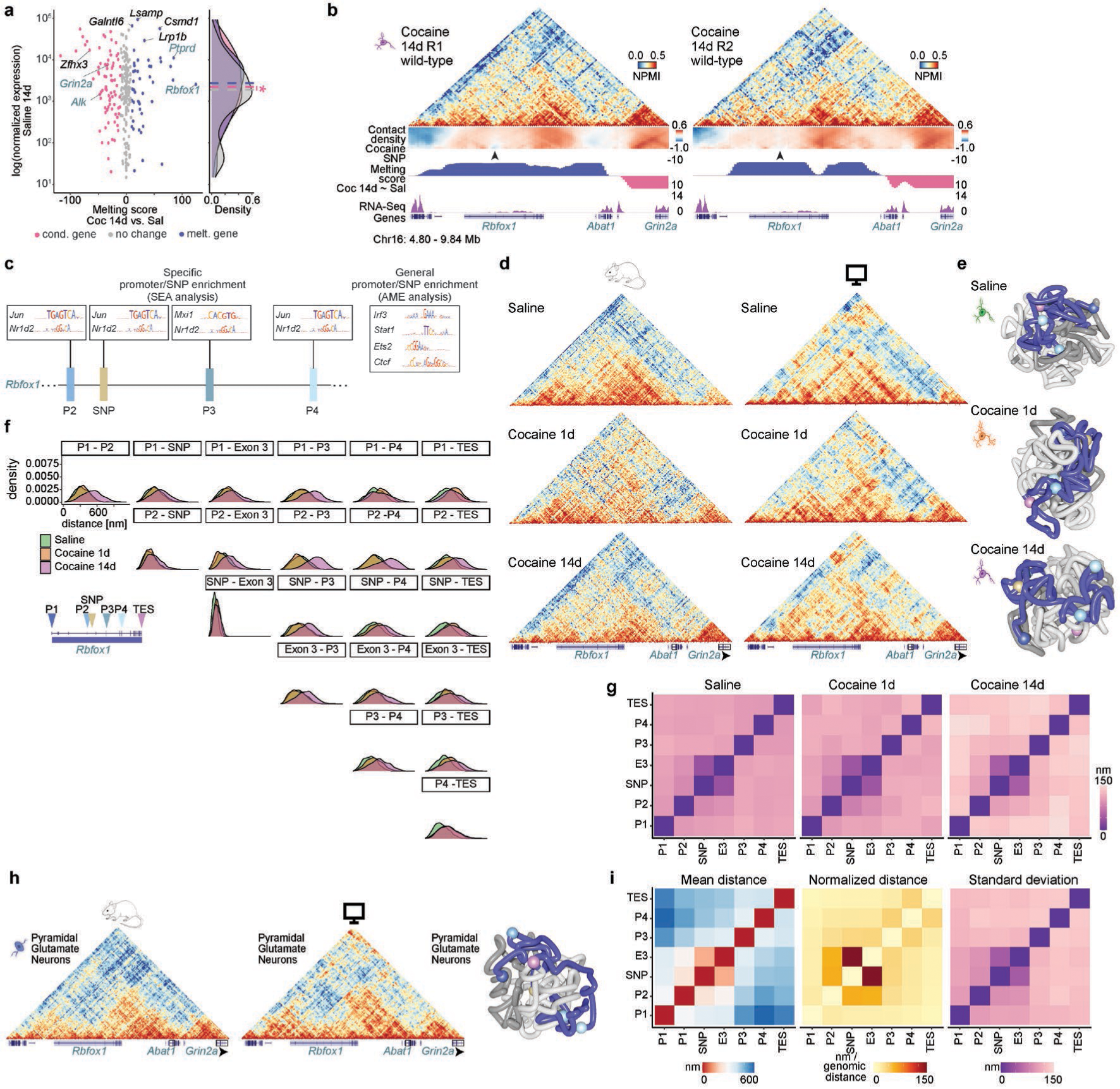
Melting of the *Rbfox1* gene becomes progressively stronger after cocaine exposure. **a,** Similar to **Fig. 3B**, but for melting scores 14 days after cocaine exposure. Condensing is associated with higher baseline (saline) transcription (one-sided Wilcoxon signed rank test, \**P* = 0.05). **b,** Log2 fold change (log2FC) distributions of melting, condensing and non-changing genes 1d after cocaine exposure. **c,** Log2FC distributions of melting, condensing and non-changing genes 14 days after cocaine exposure. n.s., not significant. For **b** and **c**, boxplot whisker length represents 1.5 times the interquartile range. **d,** Similar to **Fig. 3c** but also includes the second 14-day cocaine replicate. Melting can be seen across the gene in both GAM replicates, with the strongest melting at a putative cocaine SNP (arrowhead). **e,** Enrichment of immediate early gene and circadian TFBS motifs for the *Rbfox1* putative SNP and nearby promoters (P2, P3 and P4; enrichment +/- 500kb around the feature) using Simple Enrichment Analysis (SEA) or Analysis of Motif Enrichment (AME). Full result table can be found in **Supplementary Table 9**. **f,** Similar to **Fig. 3d** but for an additional example polymer, showing looping out of the TES and full gene decondensation 1 and 14 days after cocaine, respectively. **g,** GAM matrices from the *Rbfox1* region (chr16: 4.80 - 9.81 Mb, 30 kb resolution) compared to *in-silico* GAM reconstructed matrices from the ensemble of polymers (n=1100 for each condition) after modeling (Pearson r = 0.65, 0.56 and 0.64 for saline, cocaine 1 day and cocaine 14 days, respectively). **h,** Density estimates of pairwise distances at the *Rbfox1* gene across 1100 polymer models in saline, cocaine 1- and 14-day conditions. Multiple pairwise distances show increased contacts after cocaine exposure, most pronounced by 14 days. **i,** The variability of measured distances between the *Rbfox1* gene features are increased after cocaine exposure with highest values measured after 14 days. **j,** Similar to **Extended Data Fig. 6f, g**, but modeling from PGNs (n=1100, Pearson r=0.65, GAM data from^10^). **k,** Similar to **Fig. 3e, f** and **Extended Data Fig. 6j** showing the average distance, normalized linear distance, and distance variation between *Rbfox1* promoters, the putative SNP, SNP-proximal exon and TES for PGNs. The models, distances and variability were most similar to DNs 14 days following cocaine treatment.

**Extended Data Figure 7.**
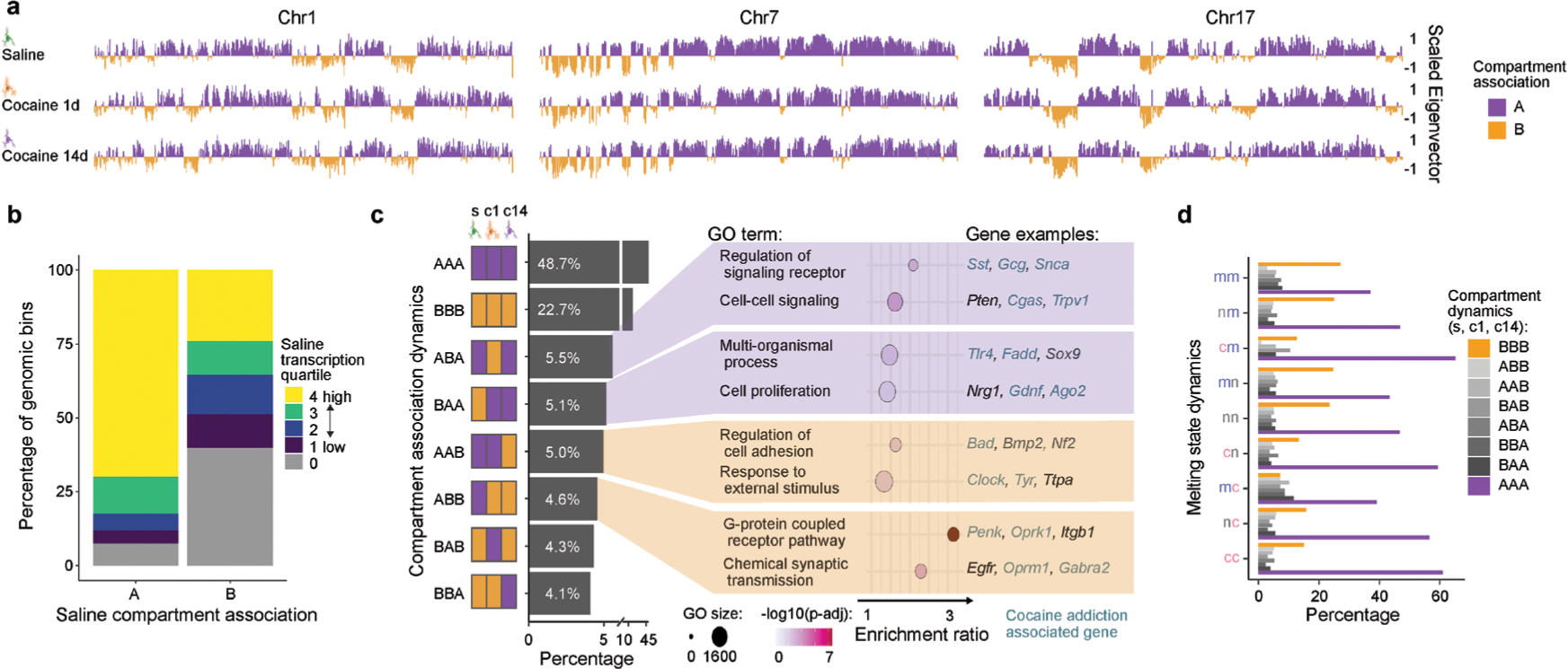
Compartment dynamics after a single cocaine exposure. **a,** Linear representation of scaled eigenvector values and compartment associations of representative chromosomes at 250 kb resolution. The full list of eigenvector values and compartment associations can be found in **Supplementary Table 8**. **b,** 67% of all genomic bins assigned to the A compartment after saline treatment contain at least one highly expressed gene, while B compartments most frequently contain non-expressed genes (42%). **c,** Frequency of compartment switches between saline, cocaine 1 and 14 day wild-type samples. Compartment regions that change following cocaine exposure (4.1 - 5.5% of bins) contain genes with functions in synaptic signalling. A complete list of gene ontology (GO) terms can be found in **Supplementary Table 10**. **d,** Most melting or condensing regions do not change compartment classification following 1 or 14 days of cocaine exposure.

**Extended Data Figure 8.**
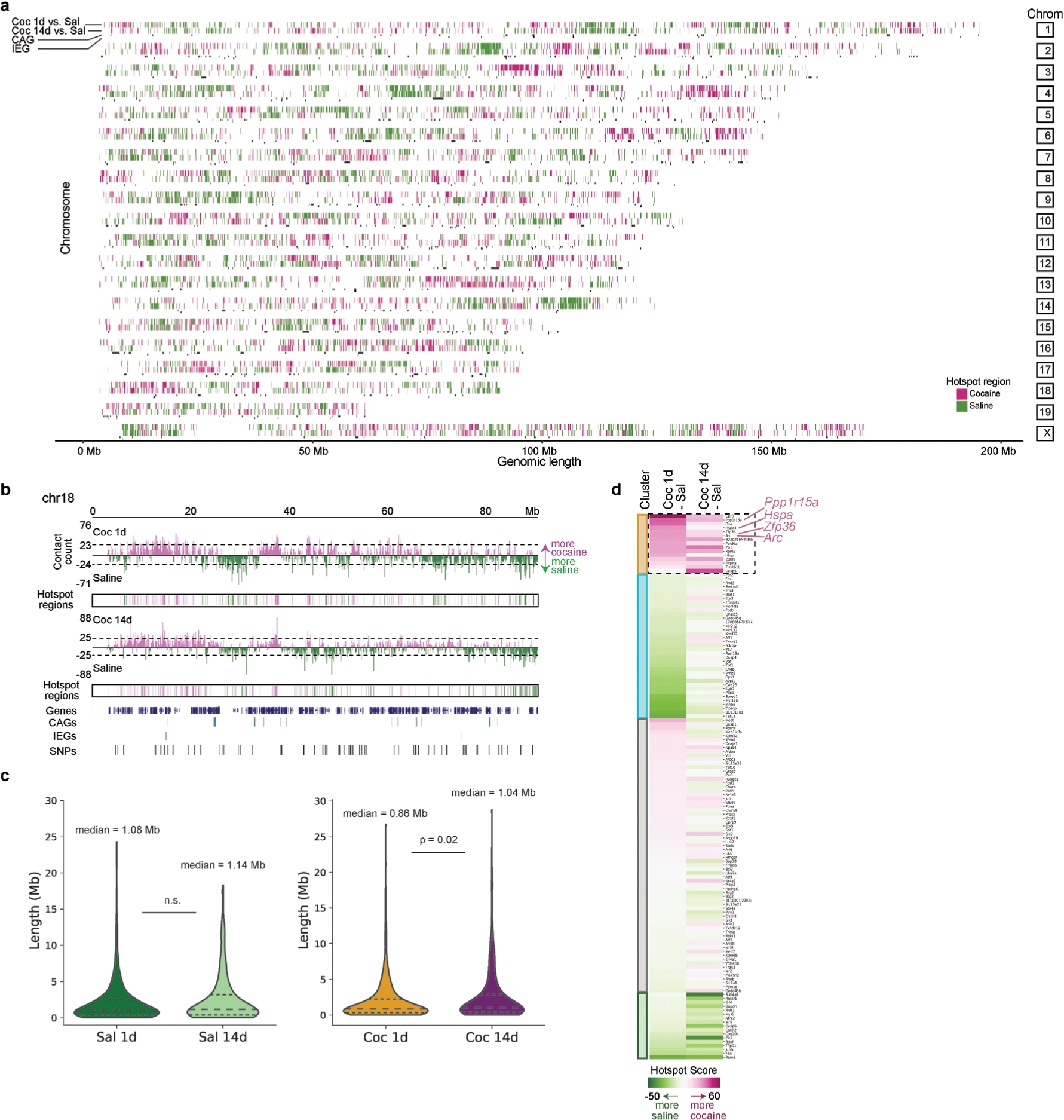
Hotspots of differential contacts are clustered in the genome and propagate during the cocaine response. **a,** Linear version of **Fig. 4c** showing cocaine (pink) and saline (green) hotspots across all chromosomes. **b,** Differential contact count per genomic window, hotspot thresholds (indicated with the dashed line), and preferred contact regions (hotspots) for chromosome 18. **c,** Genomic lengths of contiguous hotspots, calculated as the number of genomic windows containing a hotspot from a given treatment before encountering a hotspot from the other treatment. Saline hotspots were longer than 1 day cocaine hotspots (two-sided Mann Whitney U-test, *P* = 0.02), with no difference between saline and 14-day cocaine hotspots. **d,** Heatmap of hotspot scores for cocaine 1 or 14 days, compared to saline. Clustering of IEGs by hotspot score shows a group of IEGs with high hotspot scores both 1 and 14 days after cocaine. Example genes are labelled pink.

**Extended Data Figure 9.**
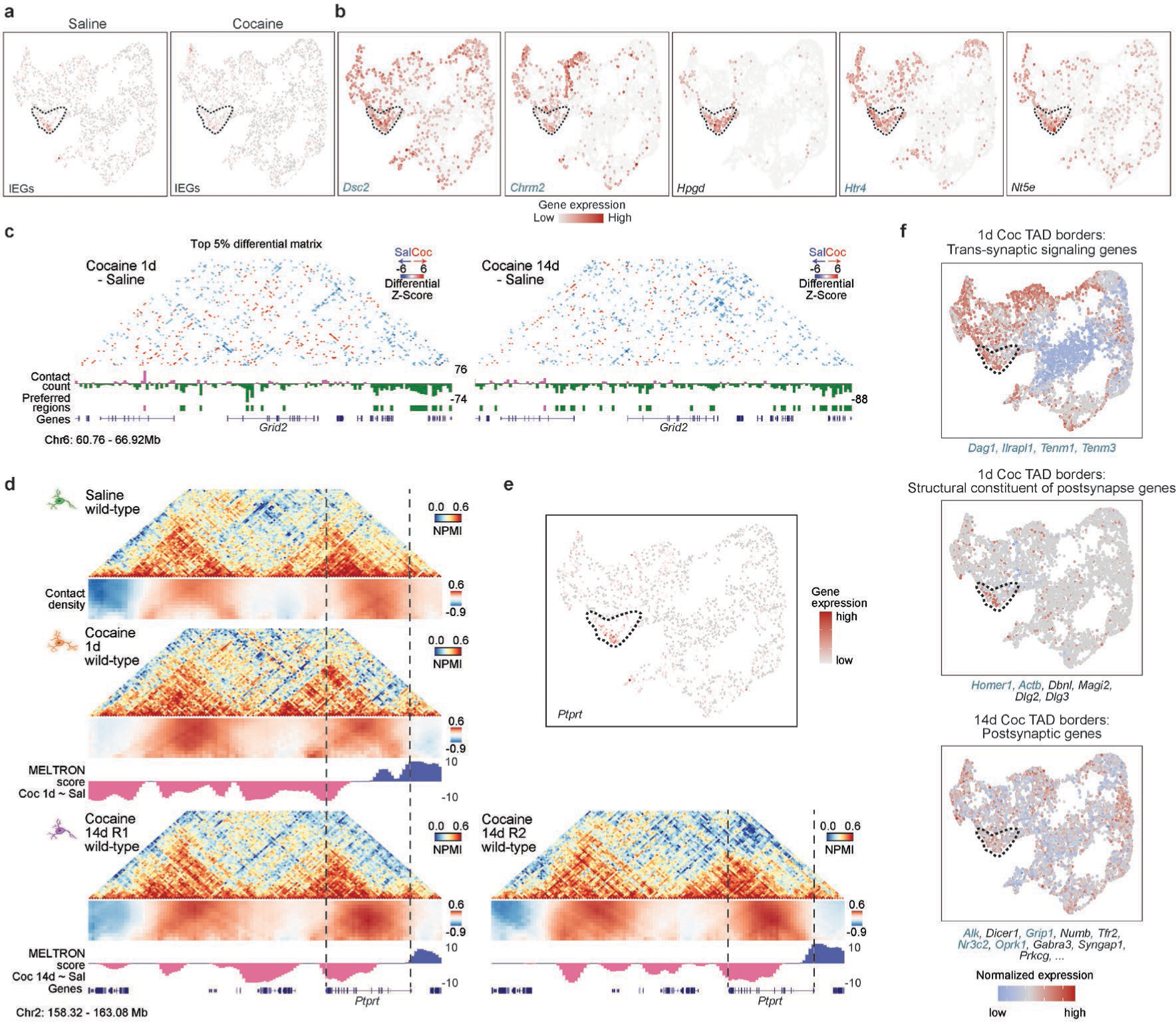
Immediate early genes and cocaine addiction associated genes are preferentially expressed in a specific DN subcluster. **a,** Similar to **Fig. 5b**, except showing the entire DN UMAP. The IEG group had the highest expression in the identified cluster (dashed lines) in saline treatment, which was lower after cocaine exposure. **b,** Related to **Fig. 5c**, example UMAPs of IEG cluster marker genes. All genes were highly expressed in the IEG cluster (dashed lines), with some marker genes expressed in other clusters. **c,** Related to **Fig. 5d**, example region containing the *Grid2* gene (chr6: 60.76 – 66.92 Mb), showing strong saline-specific contacts (differential matrix) and hotspots (tracks below matrix) compared to both 1 and 14 days after cocaine. **d,** Example region containing the *Ptprt* gene (chr2: 158.32 - 163.08 Mb), showing melting at the gene TSS, and condensing at the gene TES, both 1 and 14 days after cocaine exposure. **e,** UMAP of *Ptprt* expression shows high expression in the IEG cluster. **f,** Similar to **Fig. 5e**, except showing additional synGO gene ontology groups of genes found in cocaine-specific TAD boundaries with high expression in the IEG cluster.

**Extended Data Figure 10.**
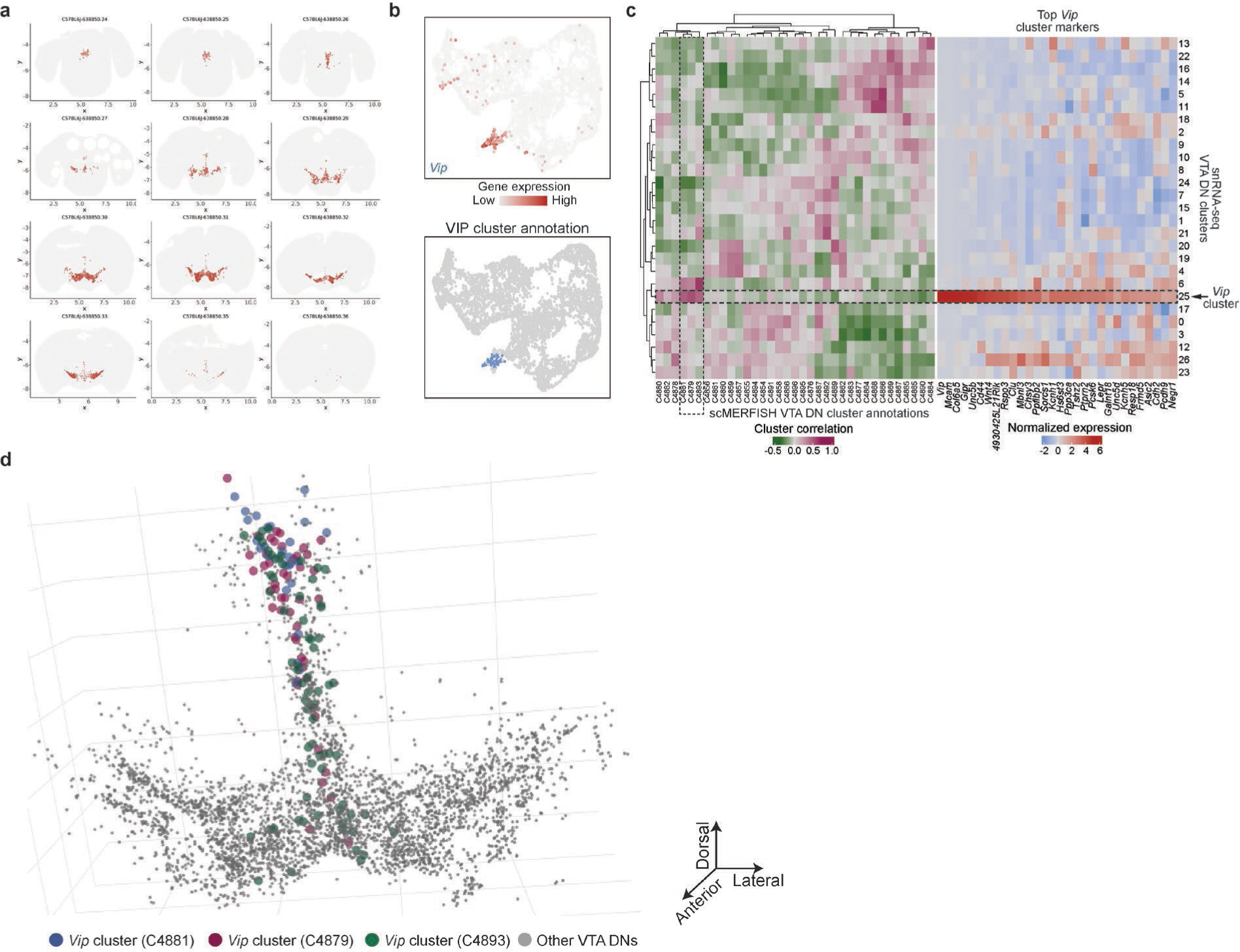
*Vip* expressing DNs are located to the VTA midline and show high dopamine auto-receptor expression. **a,** Related to **Fig. 5f**, all mouse brain slices from single-cell MERFISH (scMERFISH) spatial transcriptomic data identified to contain DNs using cell-type specific markers (*Th* expression shown). Slices are named based on their scMERFISH annotation, and range from slice 24 - 36. Note, slice 34 is not shown as none of the cells were identified as VTA DNs in the scMERFISH annotation. **b,** UMAP of *Vip* expression (top) and *Vip* cluster annotation (bottom). **c,** The scMERFISH clusters with highest correlation to the snRNA-seq *Vip* expressing cluster are C4881, C4879 and C4893. **d,** *Vip^+^* DNs (colored dots) mainly localize to the dorsal-medial VTA.

## Captions for Supplementary Videos 1 to 8

**Supplementary Video 1. *Rbfox1* example polymer 1 in saline conditions.** Rotating 3D-image of the example polymer shown in **Fig. 3d**.

**Supplementary Video 2. *Rbfox1* example polymer 2 in saline conditions.** Rotating 3D-image of the example polymer shown in **Extended Data Fig. 6f**.

**Supplementary Video 3. *Rbfox1* example polymer 1, 1 day after cocaine exposure.** Rotating 3D-image of the example polymer shown in **Fig. 3d**.

**Supplementary Video 4. *Rbfox1* example polymer 2, 1 day after cocaine exposure.** Rotating 3D-image of the example polymer shown in **Extended Data Fig. 6f**.

**Supplementary Video 5. *Rbfox1* example polymer 1, 14 days after cocaine exposure** Rotating 3D-image of the example polymer shown in **Fig. 3d**.

**Supplementary Video 6. *Rbfox1* example polymer 2, 14 days after cocaine exposure.** Rotating 3D-image of the example polymer shown in **Extended Data Fig. 6f**.

**Supplementary Video 7. IEG expressing dopamine neurons localize along the entire midline of the VTA.** Rotating 3D-image of the IEG DN cluster mapped to single-cell MERFISH data shown in **Fig. 5g**.

**Supplementary Video 8. *Vip* expressing dopamine neurons localize along the dorsal midline of the VTA.** Rotating 3D-image of the *Vip* DN cluster mapped to single-cell MERFISH data shown in **Extended Data Fig. 10d**.

## Tables

**Supplementary Table 1.** Genomic coordinates of immediate early genes (IEGs) and cocaine addiction genes (CAGs).

**Supplementary Table 2.** Marker genes of dopamine neuron subpopulations in the ventral tegmental area (VTA)

**Supplementary Table 3.** Differential gene expression table of ventral tegmental area dopamine neurons from mice treated with cocaine vs. saline treated controls.

**Supplementary Table 4.** Position of topologically associating domain (TAD) boundaries in individual samples and sample comparisons.

**Supplementary Table 5.** MELTRONIC scores and MELTRON scores of long genes.

**Supplementary Table 6.** Differential contact count and hotspot locations.

**Supplementary Table 7.** Expermiental, sequencing and QC metrics for GAM samples.

**Supplementary Table 8.** Compartment eigenvector (EV) values and A/B classification for GAM datasets.

**Supplementary Table 9.** Transcription factor binding motif enrichment results at the Rbfox1 locus.

**Supplementary Table 10.** Full list of enriched gene ontology (GO) terms and associated genes.

